# Phospholipidosis is a shared mechanism underlying the *in vitro* antiviral activity of many repurposed drugs against SARS-CoV-2

**DOI:** 10.1101/2021.03.23.436648

**Authors:** Tia A. Tummino, Veronica V. Rezelj, Benoit Fischer, Audrey Fischer, Matthew J. O’Meara, Blandine Monel, Thomas Vallet, Ziyang Zhang, Assaf Alon, Henry R. O’Donnell, Jiankun Lyu, Heiko Schadt, Kris M White, Nevan J. Krogan, Laszlo Urban, Kevan M. Shokat, Andrew C. Kruse, Adolfo García-Sastre, Olivier Schwartz, Francesca Moretti, Marco Vignuzzi, Francois Pognan, Brian K. Shoichet

## Abstract

Repurposing drugs as treatments for COVID-19 has drawn much attention. A common strategy has been to screen for established drugs, typically developed for other indications, that are antiviral in cells or organisms. Intriguingly, most of the drugs that have emerged from these campaigns, though diverse in structure, share a common physical property: cationic amphiphilicity. Provoked by the similarity of these repurposed drugs to those inducing phospholipidosis, a well-known drug side effect, we investigated phospholipidosis as a mechanism for antiviral activity. We tested 23 cationic amphiphilic drugs—including those from phenotypic screens and others that we ourselves had found—for induction of phospholipidosis in cell culture. We found that most of the repurposed drugs, which included hydroxychloroquine, azithromycin, amiodarone, and four others that have already progressed to clinical trials, induced phospholipidosis in the same concentration range as their antiviral activity; indeed, there was a strong monotonic correlation between antiviral efficacy and the magnitude of the phospholipidosis. Conversely, drugs active against the same targets that did not induce phospholipidosis were not antiviral. Phospholipidosis depends on the gross physical properties of drugs, and does not reflect specific target-based activities, rather it may be considered a confound in early drug discovery. Understanding its role in infection, and detecting its effects rapidly, will allow the community to better distinguish between drugs and lead compounds that more directly impact COVID-19 from the large proportion of molecules that manifest this confounding effect, saving much time, effort and cost.

**One Sentence Summary:** Drug-induced phospholipidosis is a single mechanism that may explain the *in vitro* efficacy of a wide-variety of therapeutics repurposed for COVID-19.

## Main Text

The outbreak of COVID-19 (Coronavirus Disease 2019) has led to multiple drug repurposing screens to find antiviral therapeutics; even a superficial survey of the literature reveals over 122 such studies over the last year(*1*). The motivation is clear—discovering and developing a new drug typically takes over a decade(*2*), while a drug previously approved for another indication can be rapidly brought to the clinic to treat an urgent threat like COVID-19. Conversely, the likelihood that a drug developed against a human disease will work against a novel virus might seem tenuous(*3*). Indeed, only a handful of these drugs are established antivirals, and these are generally restricted in activity to viruses of the same family. Thus, it has been encouraging, if not surprising, to see that over 1,974 unique drugs and investigational drugs have reported activity against SARS-CoV-2, the virus that causes COVID-19(*1*) (**Fig. 1**). Since this RNA virus encodes only 29 of its own proteins, the question of mechanism of action arises.

**Fig. 1.**
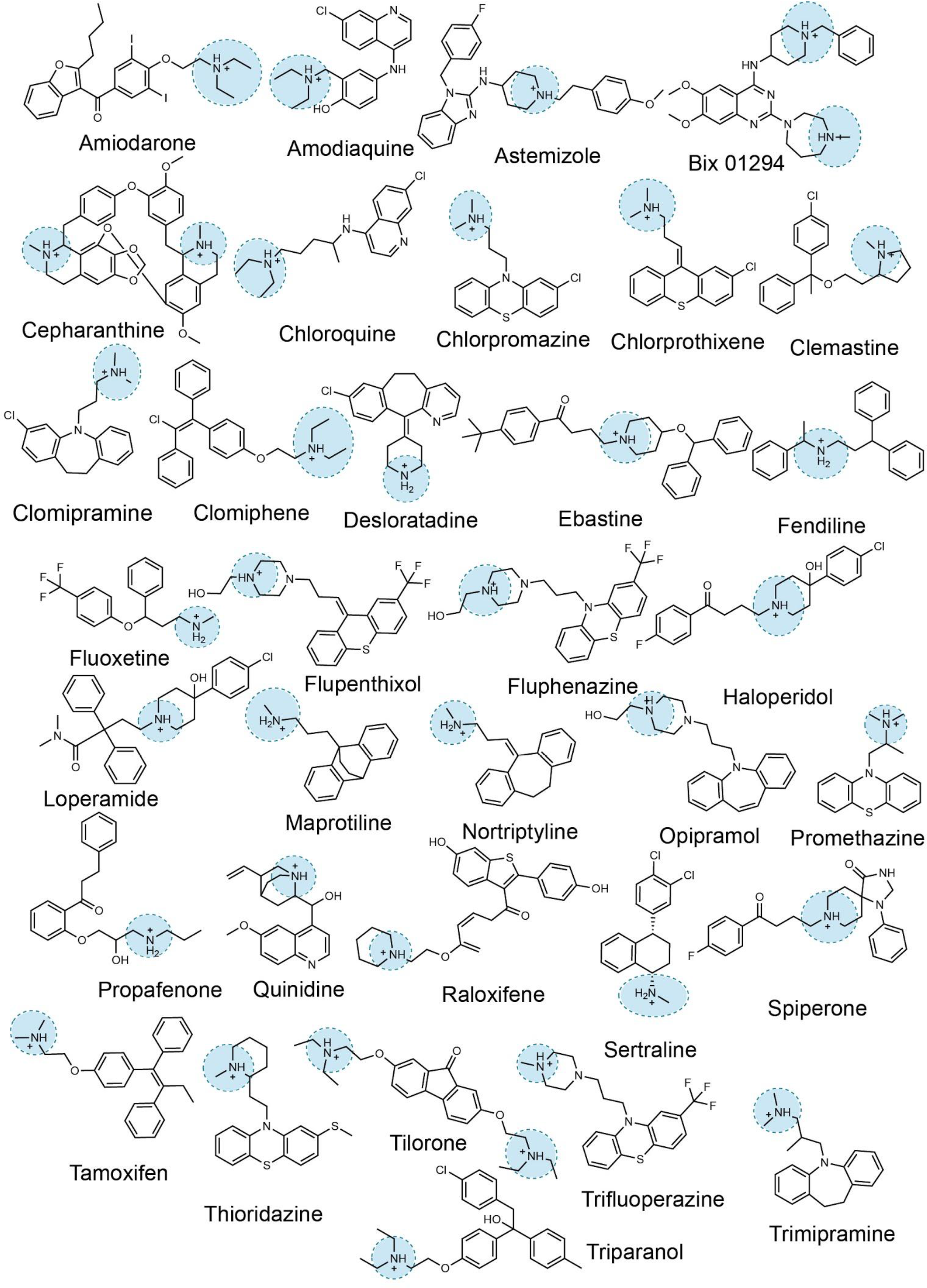
Representative examples of cationic amphiphilic drugs that are identified in SARS-CoV-2 drug repurposing screens.

Our interest in this question was motivated by our discovery that drugs binding to the sigma-1 and sigma-2 receptors were active in cells as antivirals. These receptors were identified proteomically as human proteins that interacted with the viral proteins Nsp6 and Orf9c, respectively(*4*). Encouragingly, early work showed that drugs and reagents like chloroquine, haloperidol, clemastine, and PB28—all potent against one or both sigma receptors—had cellular antiviral IC_50_ values in the 300 nM to 5 μM range. Subsequently, we investigated over 50 different molecules with a wide range of activity at these receptors. While this found molecules with relatively potent activity, structure activity relationships (SAR) found little correlation between receptor potency and antiviral efficacy (**Fig. S1**). Whereas drugs like amiodarone, sertraline, and tamoxifen had mid-to high-nM activities against viral replication, there were other sigma-active compounds, such as melperone and DTG, that were also potent on one or both sigma receptors but had no measurable antiviral effects. Intriguingly, we noticed the sigma drugs that were antiviral were all cationic at physiological pH and relatively hydrophobic, while those that were inactive against the virus were often smaller and more polar. We further noticed that this cationic-amphiphilic character was also shared among many of the hits emerging from other phenotypic screens (**Fig. 1, S10**), suggesting it was this physico-chemical property that might explain antiviral activity, instead of a specific on-target activity(*5*).

If the cationic-amphiphilic nature of this large set of molecules was itself responsible for antiviral activity, rather than the particular and widely different on-target activities of each individual drug, one would expect such a physical property to be associated with a shared cellular mechanism. Indeed, cationic amphiphilic drugs (CADs) can provoke phospholipidosis in cells and organs. This well-known side effect often encountered in late-stage drug development is characterized by the formation of lipidic vesicle-like structures in susceptible cells, and “foamy” or “whorled” membranes(*6, 7*). Though its molecular etiology remains a matter of ongoing research, drug-induced phospholipidosis (DIPL) is thought to arise by CAD disruption of lipid homeostasis. In particular, CADs are known to accumulate in intracellular compartments such as endosomes and lysosomes where they can directly or indirectly inhibit lipid processing(*6*). Importantly, modulation of these same lipid processing pathways is critical for viral replication(*8*), and inhibiting phospholipid production has previously been associated with inhibition of coronavirus replication(*9*). CADs have been previously shown to have *in vitro* activity against a range of viruses such as Severe Acute Respiratory Syndrome virus, Middle East Respiratory Syndrome virus, Ebola virus, Zika virus, Dengue virus, and filoviruses(*10*), though CAD-induction of phospholipidosis has only been proposed as a specific antiviral mechanism against Marburg virus(*11*). Finally, many of the drugs that are best-known to induce phospholipidosis, and for which it is associated with adverse events, are amiodarone(*12*) and chloroquine(*13, 14*), which we(*15*), and others (*16, 17*), had found to be potent inhibitors of SARS-CoV-2 replication *in vitro*. Other drugs from other SARS-CoV-2 phenotypic screens reported in the literature, such as chlorpromazine(*18*) and tamoxifen(*17*), had also been reported to induce phospholipidosis(*19*). Although the antiviral activity of CADs has been described, few, if any, of these CAD hits for SARS-CoV-2 have considered their phospholipidosis induction as a possible explanatory factor.

Here, we investigate the association between phospholipidosis and antiviral activity against SARS-CoV-2 in cell culture. This apparently general mechanism may be responsible for a large percentage of the drug repurposing hits reported for SARS-CoV-2, and an extraordinary amount of effort and resources lavished on drug discovery against this disease. As an effect that rarely occurs at concentrations lower than 100 nM, that does not appear to translate from *in vitro* to *in vivo* antiviral activity (as we show), and that can be a dose-limiting toxicity therapeutically(*20*), phospholipidosis may act as a confound to true antiviral drug discovery. We investigate the prevalence of this confound in SARS-CoV-2 repurposing studies, how phospholipidosis in particular correlates with inhibition of viral infection, and how to eliminate such hits rapidly so as to focus on drugs with genuine potential against COVID-19, and against other pandemics yet to arise.

## Results

### Lack of correlation between Sigma receptor activity and antiviral effect

Our studies began with an effort to find drugs that modulated either or both of the sigma-1 or the sigma-2 receptors, and were therefore expected to be antiviral against SARS-CoV-2(*4*). Fortunately, many cationic drugs, for multiple primary targets and indications, have potent off-target activities against these receptors, offering a wide field for SAR. Initial results were promising, with multiple drugs found with antiviral activity and implication of sigma-1 as a relevant antiviral target from proteomic(*21, 22*) and CRISPR(*15, 23*) screens. After testing 72 sigma ligands in cellular antiviral assays and for sigma-1/sigma-2 ligand binding, it became apparent that potency for either or both sigma receptors had little relationship to antiviral activity (**Fig. S1-S3, Table S1**), suggesting that despite sigma receptors being involved in the viral life cycle, existing sigma receptor ligands may have additional cellular effects confounding their antiviral signal. This was a matter of some consternation, since by then much effort had been lavished on this project, and some of the drugs had antiviral IC_50_s as low as 130 nM.

### Correlation of phospholipidosis and antiviral activity

Among the most potent in the antiviral assays were drugs like amiodarone, tamoxifen, and sertraline; CADs known to provoke phospholipidosis at relevant concentrations, a connection also made previously for filoviruses (*11*). Since phospholipidosis disrupts intracellular lipid homeostasis, and many viruses propagate in double membrane vesicles that may exploit the lipid pathways that are disrupted by CADs, we investigated the hypothesis that the antiviral effect we had found, and potentially found in other drug repurposing studies for COVID-19, could be explained by the drug induction of phospholipidosis.

Accordingly, we tested 19 drugs (18 CADs and 1 non-CAD known to induce phospholipidosis) for their induction of phospholipidosis in A549 cells, a human lung-derived cell line widely used for testing viral infectivity. To measure phospholipidosis, we used the well-established NBD-PE staining assay(*24*), where the vesicular lipidic bodies characteristic of the effect may be quantified by high content imaging (**Fig. 2A**). Three classes of drugs and reagents were initially investigated: **A.** Sigma-binding antiviral CADs we had discovered, like amiodarone, sertraline, chlorpromazine, and clemastine (nine total); these molecules are predicted—or known—to induce phospholipidosis; **B.** Analogs of these CADs that no longer bound sigma receptors, but were still antiviral (four total); these molecules are predicted to induce phospholipidosis despite their lack of on-target activity; and **C.** Sigma-binding, ***non***-antiviral drugs, like melperone and DTG, that were much more polar than classic CADs (two total); these molecules are predicted not to induce phospholipidosis. Of the nine sigma-binding CADs that were antiviral (class **A**), six of which were also found in the literature for COVID-19, eight induced phospholipidosis, consistent with the hypothesis (**Fig. 2A-B, S4-S5**). The only non-phospholipidosis inducing antiviral from this set was elacridar, a promiscuous P-glycoprotein inhibitor; this investigational drug may therefore be active via another mechanism. Intriguingly, CAD analogs of the potent sigma ligand PB28 that had lost their sigma-binding activity from bulky electron-withdrawing substitutions (ZZY-10-051 and ZZY-10-061, **Fig. 2B-F, S4-S7**), did induce phospholipidosis, as did the antipsychotic olanzapine and the antihistamine diphenhydramine, which are weak sigma receptor ligands but are structurally related to potent sigma receptor tricyclics (e.g., chlorpromazine) and diarylethanolamines (e.g., clemastine; class **B**). Finally, melperone and DTG, which are potent cationic sigma receptor ligands but are not antiviral, did not induce phospholipidosis (**Fig. 2A-B, S4-S5**; class **C**). These results do not prove phospholipidosis as the antiviral mechanism, but are consistent with the phospholipidosis hypothesis.

**Fig. 2.**
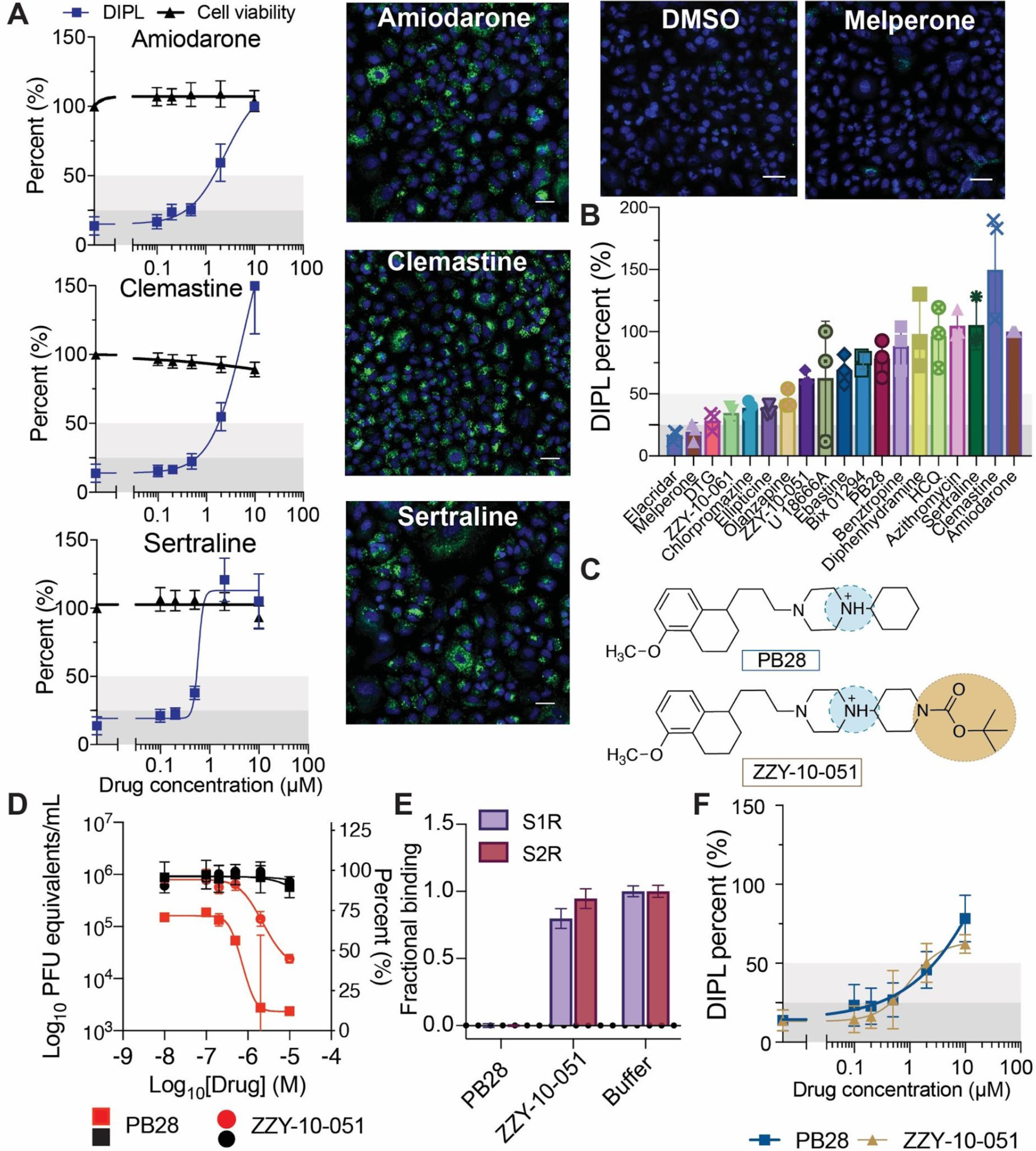
Cellular phospholipidosis may confound antiviral screening results. **A.** Examples of NBD-PE quantification of phospholipidosis in A549 cells including dose response curves. Blue = Hoechst nuclei staining, Green = NBD-PE phospholipid staining, Red = EthD-2 staining for dead cells. Scale bars = 20 µm. Amiodarone is the positive control for assay normalization; sertraline and clemastine are two examples of high phospholipidosis inducing drugs (phospholipidosis (DIPL) > 50% of amiodarone). Images of DMSO and a non-phospholipidosis inducing molecule (melperone) are included for reference. Thresholds for determining phospholipidosis power are shaded in dark grey (low phospholipidosis), light gray (medium phospholipidosis) and no shading (high phospholipidosis). **B.** Pooled DIPL amounts (mean ± SD) at the highest non-toxic concentration tested for each drug. Results were pooled from three biological and three technical replicates and were normalized amiodarone (100%) from the control wells in the same experimental batches. **C.** Structures of PB28 and its analog ZZY-10-051, the latter of which is inactive on the sigma receptors. **D**. Viral infectivity (red) and viability (black) data for PB28 (square) and ZZY-10-051 (circle) in A549-ACE2 cells. Data shown are mean ± SD from three technical replicates. **E.** Fractional binding of PB28 and ZZY-10-051 against sigma-1 (purple; S1R) and sigma-2 (maroon; S2R) normalized to a buffer control at 1.0 in a radioligand binding experiment. Data shown are mean ± SEM from three technical replicates. PB28 is a strong ligand of both sigma-1 and sigma-2 and has high displacement of the radioligands, whereas ZZY-10-051 is unable to displace the radioligands to a high degree at 1 µM. **F.** Dose response curves for PB28 (blue) and ZZY-10-051 (gold) show that these closely related analogs both induce phospholipidosis.

If phospholipidosis is responsible for CAD antiviral activity, then CADs known to induce phospholipidosis should also be antiviral, as should the rare non-CADs that are nevertheless known to induce phospholipidosis. We tested three CADs for antiviral activity, including ebastine, ellipticine, and Bix 01294, all of which are reported to induce phospholipidosis(*25*), none of which, to our knowledge, at the time of running this experiment, had been reported to inhibit SARS-CoV-2; we have since read the work of Dittmar *et al.* who also identified Bix 01294 and ebastine as drug repurposing hits against SARS-CoV-2 (*26*). We further tested the macrolide antibiotic azithromycin, also reported to induce phospholipidosis(*27*), but representing very different properties from typical CADs. We first confirmed phospholipidosis-inducing activity for these molecules, though it is important to note the difficulty of separating cytotoxicity from phospholipidosis and antiviral activity for both ellipticine and ebastine (**Fig. 2B, S4**). All four molecules were next shown to be antiviral, with IC_50_ values in the 400 nM to 3 µM range, overlapping with the activities of other CADs we and others have identified for SARS-CoV-2(*26*) (**Fig. S5**). This too was consistent with the antiviral phospholipidosis hypothesis.

For phospholipidosis to explain antiviral activity, we might expect a correlation between concentration-response curves for phospholipidosis and antiviral activity for a set of compounds. We compared phospholipidosis amounts to SARS-CoV-2 amounts for each drug individually (**Fig. 3A**). Typically, the correlations were high—not only did antiviral activity occur in the same concentration ranges where phospholipidosis occurred, but the statistically significant *R^2^* values, ranging from 0.51 to 0.94, supported a quantitative relationship between the two effects (**Fig. 3A**).

**Fig. 3.**
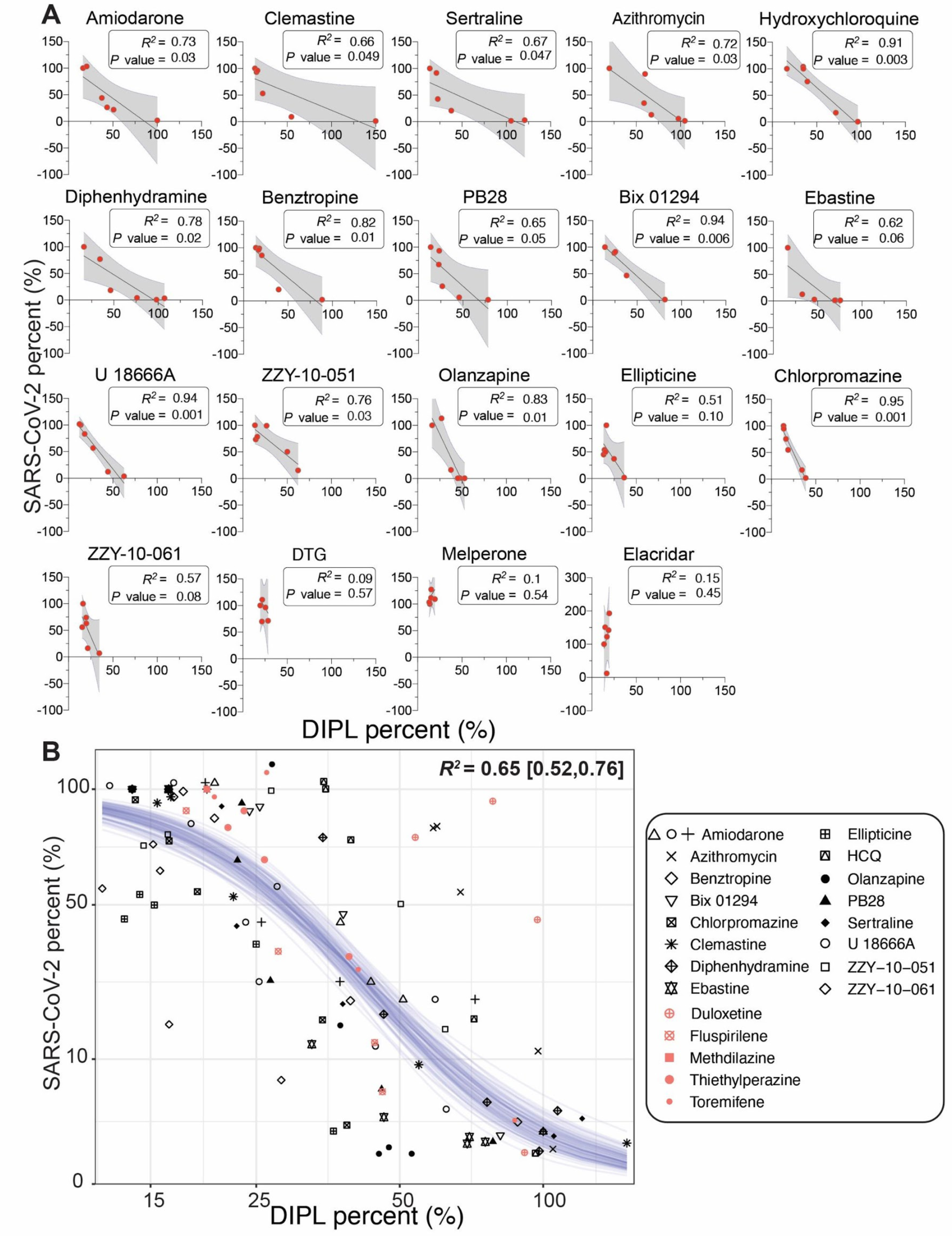
Quantitative relationship between phospholipidosis and viral amounts. **A.** Correlations between phospholipidosis (DIPL), normalized to amiodarone at 100%, and percent of SARS-CoV-2, normalized to DMSO at 100%, in the RT-qPCR assay in A549-ACE2 cells. Each dot represents the same concentration tested in both assays. A strong negative correlation emerges, with *R^2^* ≥ 0.65 and *p* ≤ 0.05 for all high and medium phospholipidosis-inducing drugs except ellipticine, which is confounded by its cytotoxicity in both experiments, ebastine, and ZZY-10-61. The latter two examples are marginally significant. **B.** The SARS-CoV-2 viral loads and induced phospholipidosis magnitude for each compound and dose in **A** are plotted as sqrt(viral_amount_mean) ∼ 10*inv_logit(hill*4/10*(log(DIPL_mean)-logIC_50_). Fitting a sigmoid Bayesian model with weakly informative priors yields parameters and 95% credible intervals of IC_50_: 43 [38, 48]%, hill: -5.6 [-7.0, -4.5], and Sigma 2.0 [0.14, 1.78]. Forty draws from the fit model are shown as blue lines. Salmon points overlaid with the model represent predicted phospholipidosis inducers from the literature (Fig. 5).

To assess if phospholipidosis could explain antiviral effect irrespective of the drug causing it, we fit a sigmoidal model through all the 107 phospholipidosis versus antiviral activity observations (comprised of six concentration measurements each for the 16 phospholipidosis-inducing drugs) and observed a strong negative relationship (*R^2^* = 0.65, 95%CI [.52, 0.76]) between induced phospholipidosis and SARS-CoV-2 viral load across all observations for all 16 drugs. Importantly, compared to the linear fit (**Fig. S8**), the sigmoid shape fit the data as well, and exhibits saturation of the effects toward the extremes. Because the biological processes of phospholipidosis and antiviral effects are saturable, the sigmoid fit was selected to represent the data.

### Concurrent measurement of viral infection and drug induced phospholipidosis

In the previous experiments, drug-induced phospholipidosis and drug antiviral activity were measured separately. To investigate how the two effects interact with each other, we next tested how viral infection affects phospholipidosis in the same cells. We dosed cells with either 1 or 10 µM of five characteristic CADs (amiodarone, sertraline, PB28, hydroxychloroquine (HCQ), and Bix 01294), followed by a mock or SARS-CoV-2 infection, and quantified phospholipidosis staining and the accumulation of viral spike protein in the same cells (**Fig. 4A, S9**). Compared to DMSO, drug treatments led to substantial increases in NBD-PE aggregates, indicating increased phospholipidosis (**Fig. S9;** 1 µM: *F* (5, 24) = 7.7, *P <* 0.001; 10 µM: *F* (5, 24) = 9.1, *P <* 0.001). At 1 µM drug concentrations, SARS-CoV-2 spike protein was readily stained, and one could visualize both spike protein and phospholipidosis staining in some of the same cells (yellow puncta), suggesting at this low concentration of drug—often close to the antiviral IC_50_ value—both phospholipidosis and viral infection were co-occuring, though even here viral staining was reduced relative to the DMSO treated controls. As drug concentration rose to 10 µM, viral spike protein staining dropped while staining for phospholipidosis increased (**Fig. S9**); there was nearly complete loss of spike protein signal with a concomitant increase in staining for phospholipidosis (**Fig. 4A**) for all treatments. To better quantify this, we repeated these experiments in seven point concentration-response for amiodarone, sertraline, and PB28. As expected, viral staining monotonically decreased as phospholipidosis increased (**Fig. 4B-C**).

**Fig. 4.**
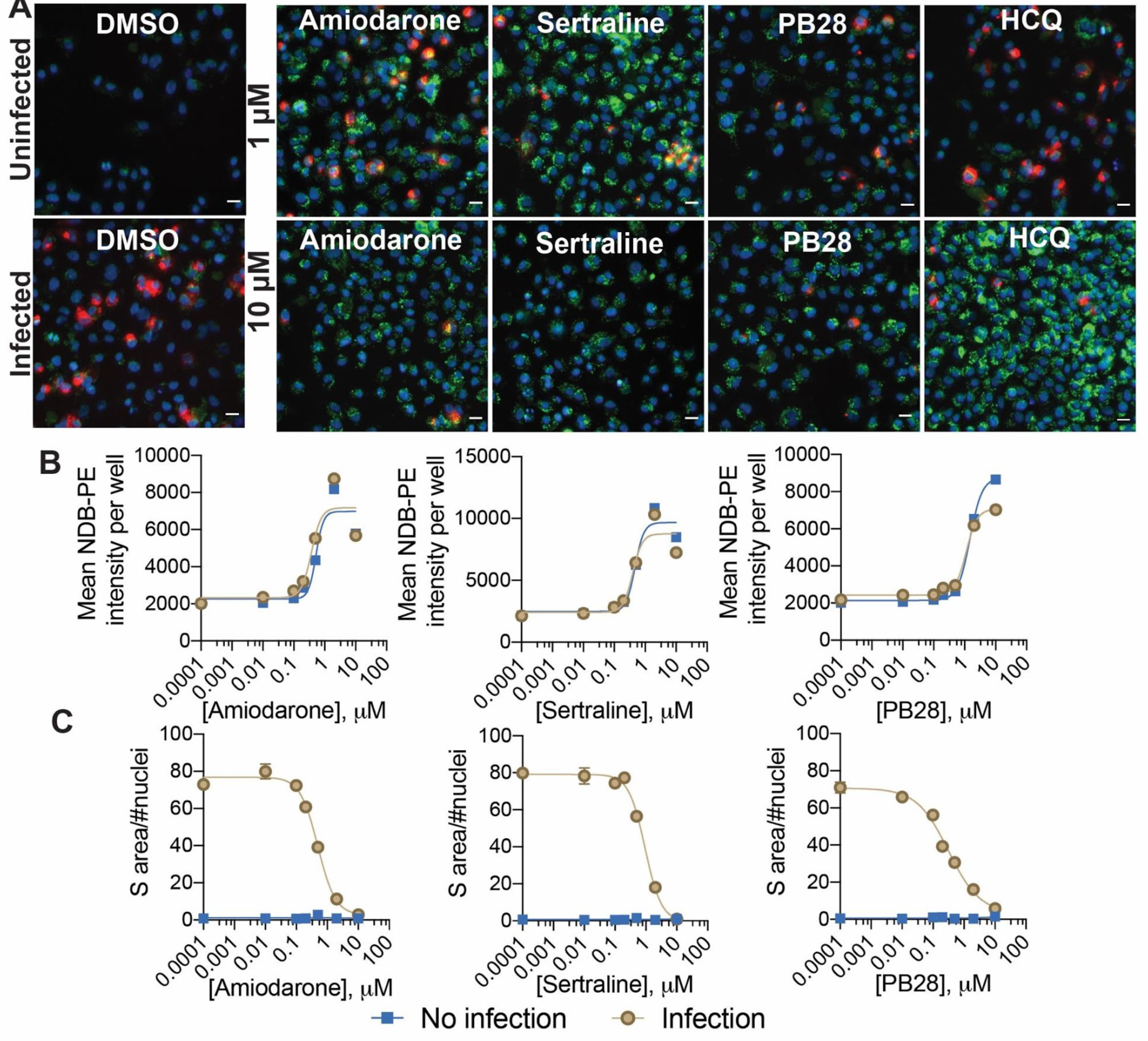
Phospholipidosis and spike protein measurements in the same cellular context. **A.** Representative images from a costaining experiment measuring phospholipidosis and SARS-CoV-2 spike protein in infected and uninfected A549-ACE2 cells. Five molecules (1 and 10 µM) and DMSO were measured; see **Fig. S9** for Bix 01294. Blue = Hoechst nuclei staining, Green = NBD-PE phospholipid staining, Red = SARS-CoV-2 spike protein staining; Yellow = coexpression of spike protein and NBD-PE. Scale bar = 20 µm. **B.** Concentration-response curves for phospholipidosis induction measured by NBD-PE staining in infected cells for three characteristic CADs. Data are mean ± SEM from four technical replicates. **C.** Spike protein in infected cells decreases as phospholipidosis increases. Data are mean ± SEM from four technical replicates.

### CADs are common among drug repurposing hits for SARS-CoV-2 and other viruses

With the strong correlation between CAD phospholipidosis and antiviral efficacy (**Fig. 3**), including drugs that have been found in multiple SARS-CoV-2 repurposing studies, such as amiodarone, sertraline, HCQ, and chlorpromazine, we wondered how prevalent phospholipidosis-inducing CADs might be in the literature. To quantify this, we investigated a subset of the 1,974 total reported repurposing hits by manually searching 12 major drug repurposing efforts for SARS-CoV-2 from the literature. These included two screens of the ReFRAME library(*28, 29*), a screen of the NCATS approved and bioactive libraries(*16*), among others (*4, 15, 17, 26, 30–34*). Together, 310 drugs, investigational drugs, and reagents were reported as active in antiviral assays against SARS-CoV-2. We used two physico-chemical features to identify likely CADs within these actives: drugs with calculated Log octanol:water coefficients above 3 (cLogP ≥ 3), and with pKa values ≥ 7.4 (*35, 36*), and then further filtered for drugs that topologically resembled known phospholipidosis inducers(*19, 25*), (using an ECFP-4-based Tanimoto coefficient (Tc) ≥ 0.4) (**Table S2**). Sixty percent of the 310 drugs surpassed the cLogP and pKa threshold; 34% were also structurally similar to a known phospholipidosis inducer (**Fig. 5A**; structures of representative Tc = 1 CADs from the literature in **Fig. 1**, and 1 > Tc ≥ 0.4 in **Fig. S10**).

**Fig. 5.**
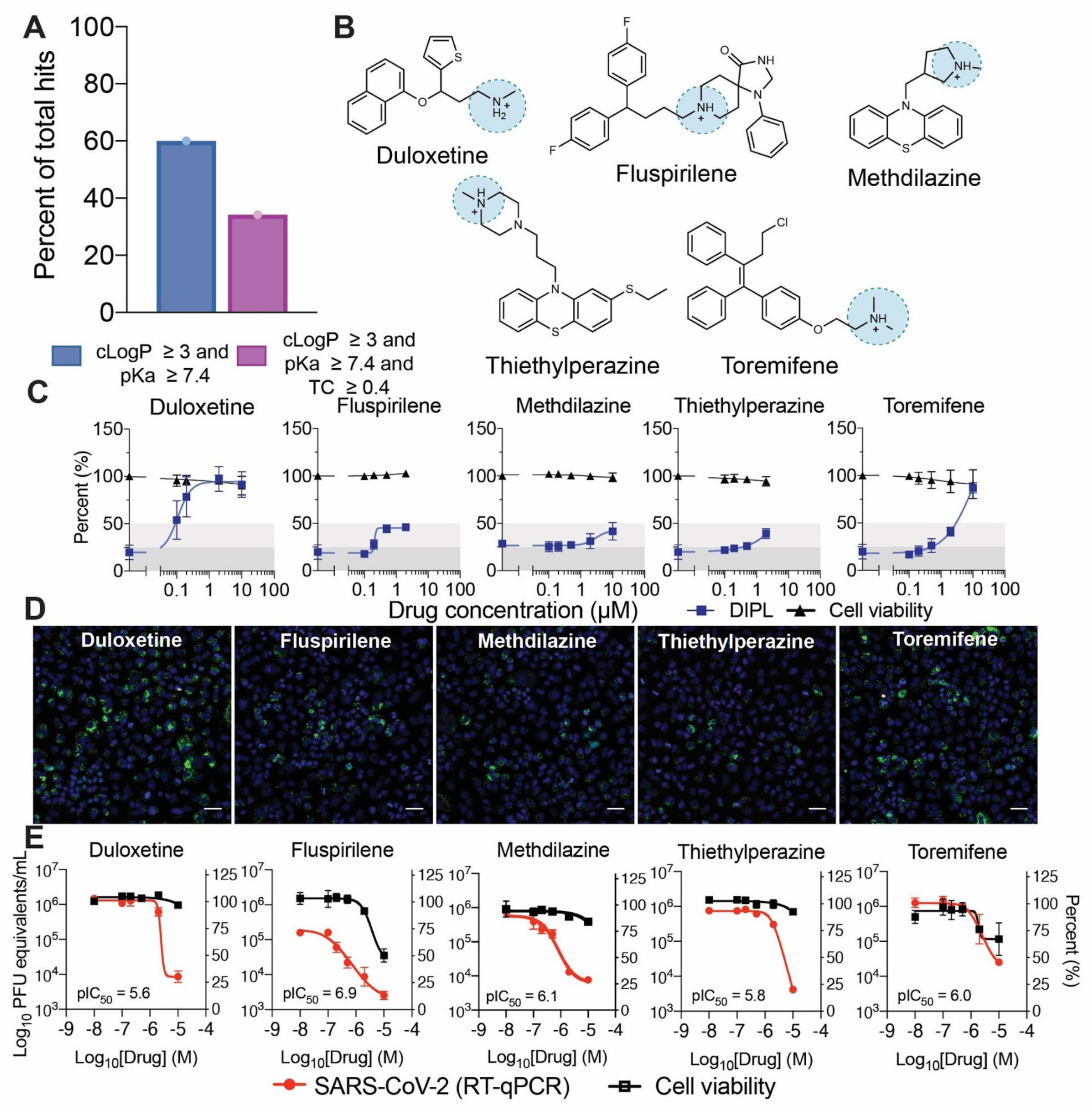
Many drugs with activity against SARS-CoV-2 are CADs that induce phospholipidosis. **A.** Percentage of total drug repurposing hits collected that pass CAD thresholds. **B.** Example repurposing hits from the literature that pass our CAD filters. **C.** Dose response curves for five predicted phospholipidosis inducers. All five induce measurable phospholipidosis (blue) with no impact on cell viability (black). **D.** Representative images of phospholipidosis quantification through NBD-PE staining in A549 cells. Blue = Hoechst nuclei staining, Green = NBD-PE phospholipid staining, Red = EthD-2 staining for dead cells. Scale bars = 20 µm. **E.** Viral infectivity (red) and cytotoxicity (black) data for five example literature CADs tested in A549-ACE2 cells. Data shown are mean ± SD from three technical replicates.

Although the two physical property filters do not capture typical phospholipidosis inducers such as azithromycin, they do capture 16 of the other 18 CADs we had tested up until now (missing only the medium phospholipidosis inducers olanzapine and ellipticine); intriguingly, nine of these, including amiodarone, sertraline, chlorpromazine, Bix 01294, clemastine, and benztropine also appeared in at least one of the 12 other repurposing studies. To probe the reliability of this association, we tested another five drugs that passed our filters, and had been reported as antiviral against SARS-CoV-2, for their ability to induce phospholipidosis (**Fig. 5B**). All five were active in the NBD-PE assay to varying extents (**Fig. 5C**). For example, duloxetine and toremifene were strong phospholipidosis inducers (> 50% of the strong phospholipidosis inducer amiodarone), while fluspirilene, methdilazine, and thiethylperazine were medium inducers (50% > DIPL > 25% of amiodarone). We were able to confirm SARS-CoV-2 antiviral activity for these drugs in our own hands (**Fig. 5D**). Additionally, these molecules fit into the sigmoidal model relating phospholipidosis amount to reduction in viral load described above (salmon points overlaid with sigmoidal model; **Fig. 3B**). Finally, we note a preliminary identification of 30 CADs, 19 of which overlap with the literature-derived SARS-CoV-2 list, active against other viruses including Middle East Respiratory virus and Severe Acute Respiratory virus(*37*), Ebola virus(*38–40*), Marburg virus(*40, 41*), Hepatitis C virus(*42*), and Dengue virus(*43*) (**Table S3**). It may be that most drugs identified in antiviral repurposing assays against SARS-CoV-2 and other viruses are CADs whose antiviral activities can be attributed to a phospholipidosis mechanism.

### Animal efficacy for phospholipidosis-inducing and non-phospholipidosis-inducing repurposed drugs

Though phospholipidosis is considered a drug-induced side effect, it remains possible that the effect can be leveraged for antiviral efficacy. Accordingly, we tested four of the repurposed drugs most potent against SARS-CoV-2 *in vitro,* whose activity is seemingly explained by phospholipidosis: amiodarone, sertraline, PB28 (three molecules tested above) and tamoxifen (a drug often reported in the literature to induce phospholipidosis)(*6, 19*), for efficacy in a murine model of COVID-19(*44*). In the same model, we also tested two antiviral drugs that do not induce phospholipidosis: remdesivir, a drug that may be considered a success of repurposing, and elacridar, an antiviral compound that was not a phospholipidosis inducer (**Fig. 2B**). In pharmacokinetic studies, all molecules had relatively long half-lives, especially in the lung where tissue C_max_ values often exceeded 10 μM after a 10 mg/kg dose; this C_max_ was 10 to 1000 times higher than the *in vitro* efficacy of these drugs against SARS-CoV-2, suggesting that exposure would be high enough for plausible efficacy (**Table S4-S8**). Guided by the pharmacokinetic parameters of each drug, mice were dosed either once per day (amiodarone and elacridar) or twice per day (remdesivir, PB28, tamoxifen, and sertraline), for a total of three days (see **Materials and Methods**). Two hours following the first dose, mice were intranasally infected with 1 × 10^4^ PFU of SARS-CoV-2, followed by viral titer quantification at the end of the three-day infection period. Notwithstanding their high lung exposure, the four phospholipidosis-inducing drugs—amiodarone, sertraline, PB28, and tamoxifen had no substantial effect on viral propagation in the mice. Conversely, the well-known antiviral remdesivir reduced viral load by two to three orders of magnitude in the mice, while the cationic drug elacridar, which had shown antiviral activity without phospholipidosis, also showed a modest antiviral effect (**Fig. 6**). Unfortunately, mice dosed higher than 3 mg/kg with elacridar exhibited toxicities that limited further study. Taken together, despite the high lung exposure of these drugs, their *in vitro* activities do not translate to *in vivo* action, at least at the tested conditions. These observations suggest that cationic amphiphilic drugs whose antiviral activity arises due to phospholipidosis may not be viable candidates for clinical progression. We do not rule out *purposefully* targeting phospholipidosis as an antiviral mechanism, though doing so may require a target-based, rather than a physical property-based approach.

**Fig. 6.**
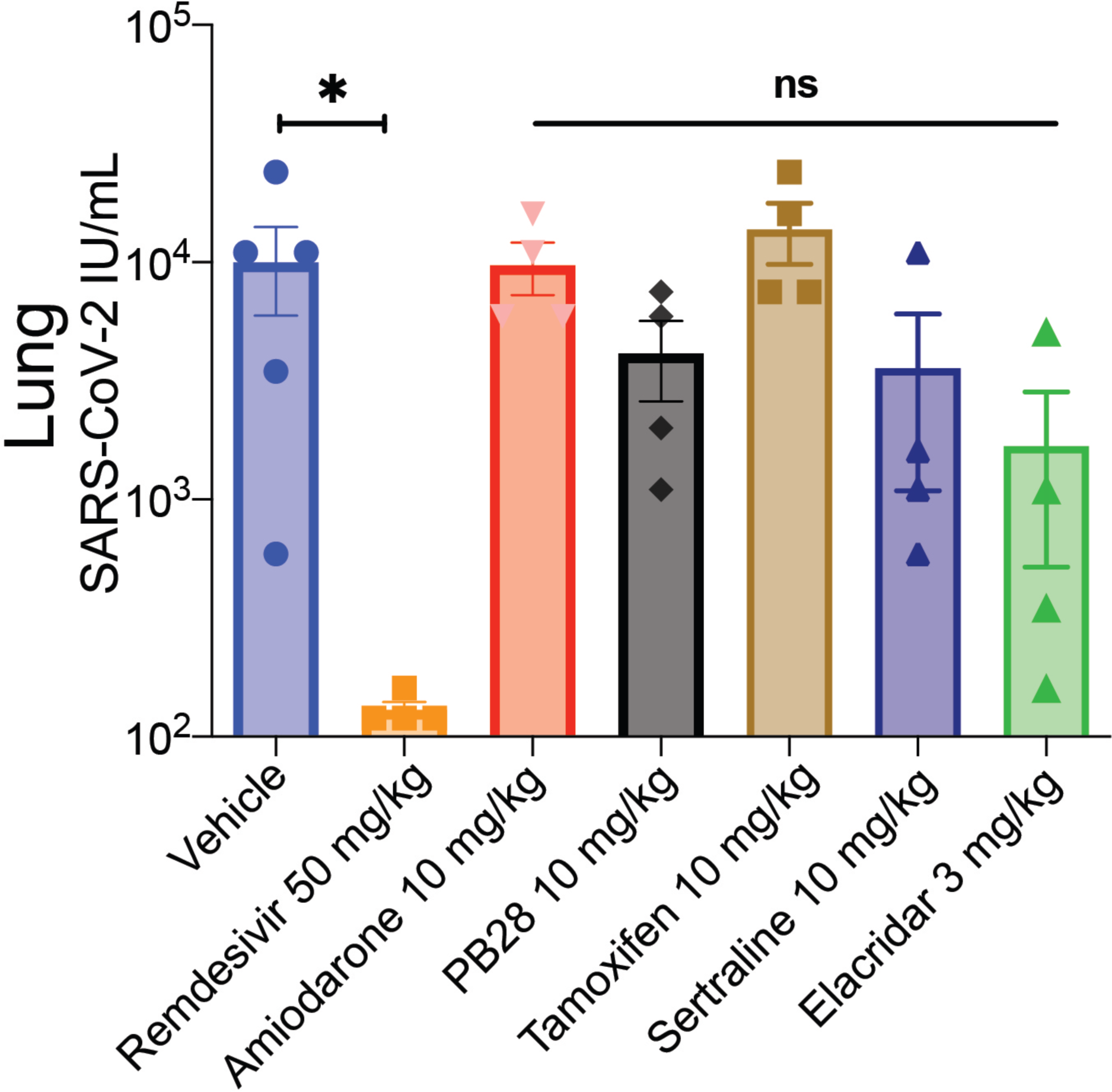
In murine viral replication, four phospholipidosis inducing drugs are not efficacious *in vivo.* Six different drugs were dosed to mice for three days with a two-hour preincubation before SARS-CoV-2 treatment. Lung viral titers were quantified. Only remdesivir, positive control, induced significant decreases in viral load (vehicle *N* = 5; remdesivir *N* = 4, \**P* < 0.05). The other five drugs, four of which cause phospholipidosis, did not significantly affect viral titers.

## Discussion

The emergence of COVID-19 has motivated an intense effort to repurpose drugs as SARS-CoV-2 antivirals. From these studies, an extraordinary number of potentially promising hits have emerged(*1*). A key observation from this work is that many, perhaps most, of these repurposed drugs are active in antiviral assays because they are cationic amphiphiles that induce phospholipidosis (**Fig. 1, 5, S10**). Thus, despite their diverse chemotypes, a single mechanism of action may explain cellular antiviral activity of these drugs. Phospholipidosis disrupts lysosomal lipid catabolism and trafficking, as reflected in the formation of the pathological vesicular structures that characterize it. This disruption may underlie the antiviral effects of these repurposed drugs, perhaps through effects on the double membrane vesicles that the virus itself creates and on which it depends. Quantitatively, there is a close *in vitro* correlation between drug-induced phospholipidosis and antiviral activity, both drug-by-drug and over the set of drugs tested here (**Fig. 3**). The effect is predictive: molecules that induce phospholipidosis are antiviral over the same concentration range, irrespective of whether they are CADs or not (e.g., azithromycin), while molecules that are related by target activity to the CADs, but are more polar and do not induce phospholipidosis (e.g., melperone and DTG), are not antiviral. Correspondingly, one can modify an antiviral drug that acts on a particular target, like the canonical sigma receptor ligand PB28, so that it loses measurable target binding, but as long as it keeps its CAD character it retains antiviral activity (e.g., ZZY-10-051, **Fig. 2D**). Unfortunately, CAD induction of phospholipidosis, at least at the potencies observed here, does not appear to translate to *in vivo* antiviral activity in the tested conditions (**Fig. 6**). More encouragingly, this study illuminates a method to rapidly identify drugs that are acting as confounds in cellular antiviral screens, allowing one to eliminate them from further study and effort and to rapidly focus on other drugs with true potential.

Although the molecular mechanisms for the antiviral effects of phospholipidosis remain unclear, certain associations may be tentatively advanced. SARS-CoV-2, like many viruses, subverts the cell to produce double membrane vesicles in which it replicates(*45–47*)—these vesicles provide sheltered environments not only for reproduction but also for evasion of innate immunity(*48, 49*). Disruption of lipid homeostasis by the induction of phospholipidosis may lead to the formation of lipid structures that compete with or disrupt the formation of viral double membrane vesicles, reducing viral replication. Alternatively, the disruption of lysosomal(*50*) and endosomal(*51*) compartments by aggregating phospholipids and CAD-induced transient shifts in compartmental pH(*52*) may further affect viral entry and propagation(*53*). For example, it is possible that alterations to these compartments disrupt cathepsins that have been shown to prime the SARS-CoV-2 spike protein facilitating viral entry(*54*). For all these reasons, targeting the endosomal-lysosomal pathway has been suggested as a valuable strategy against SARS-CoV-2 infection(*55*), but developing potent and targeted inhibitors remains challenging. Of course, these mechanisms remain unproven, and currently are supported only by correlation, but they suggest a route for further research.

The cost to the community of investments in what appears to be a confound needs to be considered as a guidance for strategy in future pandemics. According to the DrugBank(*56*) COVID-19 dashboard, which collates data from the U.S. National Library of Medicine’s ClinicalTrials.gov and the World Health Organization’s International Clinical Trials Registry Platform, CADs reported to have antiviral activity have been promoted into an astonishing 316 Phase I to Phase III clinical trials against COVID-19 (**Table S9**). Admittedly, 57% of these trials (180 of the 316) solely study the phospholipidosis-inducing CADs hydroxychloroquine (**Fig. 3A**, top row) or chloroquine, both of which already failed to show efficacy in varying cell types(*57*), rodent models(*58*), and clinical trials(*59*), but that still leaves 136 trials across 33 other predicted or known phospholipidosis-inducing CADs. Taking conservative estimates of what general anti-infective trials cost, from a 2014 US Department of Health and Human Services report(*60, 61*), we estimate the cost of the clinical trials component alone, over the last year, for phospholipidosis-inducing CADs to be over $6 billion US dollars (**Table S9**).

Certain caveats merit airing. **First**, the correlation we observe between antiviral activity and phospholipidosis, as strong as it is, does not illuminate the mechanism by which phospholipidosis is itself antiviral. Phospholipidosis itself is only partly understood mechanistically, and there are no known genetic or chemical ways to inhibit drug-induced phospholipidosis, nor are there reliable target-selective reagents to induce it. **Second**, there is no consistent standard in the field as to the physical properties that will predict whether a molecule will induce phospholipidosis, and there are even non-CAD molecules that induce it (e.g., azithromycin). Thus, we have chosen conservative criteria to predict phospholipidosis-inducing CADs; while we believe that these will have relatively few false positive predictions, many phospholipidosis-inducing drugs may be missed. **Third**, phospholipidosis is a confound that only affects drugs repurposed for direct antiviral activity—it is irrelevant for drugs like dexamethasone (*62*) and for the CAD fluvoxamine (*63*) which have been repurposed for immunomodulatory treatment of COVID-19, nor is it relevant for antiviral CADs whose activity against the virus is well-below the range where phospholipidosis occurs. **Fourth**, our estimates of the clinical trial costs of CAD advancement for COVID-19 are clearly rough. If we have inadvertently included CADs advanced for immunomodulatory treatment, for instance, they will be too high. This would be balanced by the conservative numbers that we have used to estimate trial costs, and by our conservative method of predicting molecules that are phospholipidosis-inducing CADs (of the 35 drugs identified in our analysis, 12 have been directly shown to induce phospholipidosis). Finally, we do not exclude that phospholipidosis itself might be exploited for genuinely therapeutic antiviral effect, though we suspect that will have to go through a different, presumably more target-directed mechanism than that exploited by the CADs studied here.

These caveats should not obscure the central observation of this study. Many drugs repurposed for antiviral activity against SARS-CoV-2 are cationic amphiphiles, and despite their diverse structures and multiple targets, many likely have their antiviral effects via a single shared mechanism: phospholipidosis. Both because of the adverse drug reactions with which it is associated, and the limited efficacy to which it leads—rarely better than 100 nM—drugs active due to phospholipidosis are unlikely to translate to therapeutic efficacy *in vivo* (**Fig. 6**). What this study teaches is a rapid way to distinguish drug confounds arising from phospholipidosis, on which tremendous resources have been lavished over the last year, from the genuinely effective drugs—such as the antiviral remdesivir(*64*), and the immunomodulators dexamethasone(*62*), baricitinib(*65*) —that are so desperately needed. Doing so will be important not only for COVID-19, but also for the future pandemics that we can expect to emerge.

## Terminology

COVID-19: Coronavirus Disease 2019
SAR: structure-activity relationship
CAD: cationic amphiphilic drug
DIPL: drug-induced phospholipidosis
HCQ: hydroxychloroquine
Tc: Tanimoto coefficient
EthD-2: Ethidium homodimer-2

## Acknowledgements

We gratefully acknowledge the Région Ile-de-France (program DIM1Health) for the use of the Institut Pasteur imaging facility. We thank N. Aulner for assistance with image requisition and A. Danckaert for assistance with image analysis. We thank T.R. O’Meara for reading the manuscript. We thank K. Obernier, M. Bouhaddou, & J.M. Fabius for contributions to the overall COVID-19 effort at QCRG.

## Funding

Supported by grants from the Defense Advanced Research Projects Agency (HR0011-19-2-0020 to B.K.S., N.J.K., A.G.-S., and K.M.S.); NIGMS R35GM122481 (to B.K.S.); National Institutes of Health (P50AI150476, U19AI135990, U19AI135972, R01AI143292, R01AI120694, P01AI063302, and R01AI122747 to N.J.K.); Excellence in Research Award (ERA) from the Laboratory for Genomics Research (LGR to N.J.K.); a collaboration between UCSF, UCB, and GSK (#133122P to N.J.K.); a Fast Grant for COVID-19 from the Emergent Ventures program at the Mercatus Center of George Mason University (to N.J.K.); the Roddenberry Foundation (to N.J.K.); funding from F. Hoffmann-La Roche and Vir Biotechnology; gifts from QCRG philanthropic donors (to N.J.K.); funding from Institut Pasteur (to O.S.); Urgence COVID-19 Fundraising Campaign of Institut Pasteur (to O.S.); Labex IBEID (ANR-10-LABX-62-IBEID, to O.S.); ANR/FRM Flash Covid PROTEO-SARS-CoV-2 (to O.S.); IDISCOVR (to O.S.); National Institutes of Health (R01GM119185 to A.C.K.); and Z.Z. is a Damon Runyon fellow supported by the Damon Runyon Cancer Research Foundation (DRG-2281-17).

## Author Contributions

The following authors designed and conceptualized the study: T.A.T, V.V.R, M.J.O., Z.Z., K.M.S., F.M., M.V., F.P., & B.K.S. The following authors performed experiments or data acquisition: V.V.R., B.F., A.F., B.M., T.V., Z.Z., A.A., H.R.O., & K.M.W. The following authors conducted formal data analysis: T.A.T, V.V.R, B.F., A.F., M.J.O., B.M., T.V., Z.Z., A.A., J.L., H.S., K.M.W., & F.M. The following authors supervised or managed research: M.J.O., K.M.W., N.K., L.U., K.M.S., A.C.K., AG.-S., O.S., F.M., M.V., F.P., & B.K.S. The following authors drafted the original manuscript: T.A.T., V.V.R., M.J.O, B.M., Z.Z., K.M.W., M.V., F.P., & B.K.S. All authors reviewed the manuscript prior to submission.

## Competing interests

B.F., A.F., H.S., L.U., F.M, & F.P are employees of Novartis. A.C.K. is a founder and advisor to Tectonic Therapeutic, Inc., and the Institute for Protein Innovation. K.M.S. has consulting agreements for the following companies, which involve monetary and/or stock compensation: Black Diamond Therapeutics, BridGene Biosciences, Denali Therapeutics, Dice Molecules, eFFECTOR Therapeutics, Erasca, Genentech/Roche, Janssen Pharmaceuticals, Kumquat Biosciences, Kura Oncology, Merck, Mitokinin, Petra Pharma, Revolution Medicines, Type6 Therapeutics, Venthera, Wellspring Biosciences. The Krogan Laboratory has received research support from Vir Biotechnology and F. Hoffmann-La Roche. N.J.K. has consulting agreements with Maze Therapeutics and Interline Therapeutics and is a shareholder of Tenaya Therapeutics. The García-Sastre Laboratory has received research support from Pfizer, Senhwa Biosciences, and 7Hills Pharma; and A.G.-S. has consulting agreements for the following companies involving cash and/or stock: Vivaldi Biosciences, Contrafect, 7Hills Pharma, Avimex, Valneva, Accurius, and Esperovax.

## Data and materials availability

All data is available in the manuscript or the supplementary materials. Further information and requests for resources and reagents should be directed to B.K.S., F. P., M.V, & F.M.

## Supplementary Material

### Materials and Methods

#### Competition Binding Assays

Competition curves were performed using membranes from Expi293F cells (Thermo) transiently overexpressing either the human sigma-1 or the human sigma-2 receptors. For binding assays, ^3^H-(+)-pentazocine and ^3^H-DTG were used as the radioactive probes for sigma-1 and sigma-2, respectively. Membranes were incubated in a 100 μL reaction with 50 mM Tris (pH 8.0), 0.1% (w/v) bovine serum albumin, 10 nM radioligand, and eight concentrations of the competing cold ligand. Reactions were incubated for two hours at 37 °C and then were terminated by filtration through a glass fiber filter using a Brandel harvester. Glass fiber filters were pre-soaked in 0.3% (v/v) polyethylenimine for 30 minutes at room temperature before harvesting. All reactions were performed in triplicate using a 96-well block format. After the membranes were transferred to the filters and washed, the filters were soaked in 5 mL Cytoscint scintillation fluid (MP Biomedicals) overnight, and radioactivity was measured using a Beckman Coulter LS 6500 scintillation counter. K_D_ values for each receptor were calculated in GraphPad Prism (San Diego, CA) using a saturation binding assay with eight concentrations of the radioactive ligand. Non-specific binding was measured in the presence of 10 μM haloperidol. The K_D_ of the sigma-1 radioligand was measured to be 21 nM and K_D_ of the sigma-2 radioligand was measured to be 15 nM, and these values were used to calculate K_i_ values for cold ligands in GraphPad Prism.

#### Antiviral drug and cell viability assays at Institut Pasteur

A549-ACE2 cells were propagated at 37°C, 5% CO2 in DMEM supplemented with 10% FBS and 20 μg/mL blasticidin S. 6250 of these cells per well were seeded into 384-well plates in DMEM (10% FBS) and incubated for 24 hours at 37°C, 5% CO2. Two hours prior to infection, the media was replaced with 50 μL of DMEM (2% FBS) containing the compound of interest at the indicated concentration. At the time of infection, the media was replaced with virus inoculum (multiplicity of infection, MOI = 0.1 PFU/cell) and incubated for one hour at 37°C, 5% CO2. The SARS-CoV-2 strain used (BetaCoV/France/IDF0372/2020) was propagated once in Vero-E6 cells and is a kind gift from the National Reference Centre for Respiratory Viruses at Institut Pasteur, Paris, originally supplied through the European Virus Archive goes Global platform. Following the adsorption period, the inoculum was removed, replaced with 50 μL of drug-containing media, and cells were incubated for an additional 72 hours at 37°C, 5% CO2. At this point, the cell culture supernatant was harvested and viral load was assessed by RT-qPCR. Briefly, the cell culture supernatant was collected, heat inactivated at 95°C for 5 minutes and used for RT-qPCR analysis. SARS-CoV-2 specific primers targeting the N gene region: 5′-TAATCAGACAAGGAACTGATTA-3′ (Forward) and 5′-CGAAGGTGTGACTTCCATG-3′ (Reverse) were used with the Luna Universal One-Step RT-qPCR Kit (New England Biolabs) in an Applied Biosystems QuantStudio 6 thermocycler, with the following cycling conditions: 55°C for 10 min, 95°C for 1 minute, and 40 cycles of 95°C for 10 seconds, followed by 60°C for 1 minute. The number of viral genomes is expressed as PFU equivalents/mL, and was calculated by performing a standard curve with RNA derived from a viral stock with a known viral titer. Data was fit using nonlinear regression and IC_50_s for each experiment were determined using GraphPad Prism software.

Cell viability was assayed in uninfected, drug-treated cells using the CellTiter-Glo assay following the manufacturer’s instructions (Promega). Luminescence was measured in a Tecan Infinity 2000 plate reader, and percentage viability calculated relative to untreated cells (100% viability) and cells lysed with 20% ethanol (0% viability), included in each experiment.

#### Cells and viruses for Anti-NP Immunofluorescence

Vero E6 (ATCC, CRL-1586) cells were maintained in DMEM (Corning) supplemented with 10% FB (Peak Serum) and Penicillin/Streptomycin (Corning) at 37°C and 5% CO2. This cell line was regularly screened for mycoplasma contamination using the Universal Detection Kit (ATCC, 30-1012K). Cells were infected with SARS-CoV-2, isolate USA-WA1/2020 (BEI Resources NR-52281) under biosafety level 3 (BSL3) containment in accordance to the biosafety protocols developed by the Icahn School of Medicine at Mount Sinai. Viral stocks were grown in Vero E6 cells as previously described(*66*) and were validated by genome sequencing.

#### Antiviral drug and cell viability assays at Mt. Sinai

Two thousand Vero E6 cells were seeded into 96-well plates in DMEM (10% FBS) and incubated for 24 hours at 37°C, 5% CO2. Two hours before infection, the medium was replaced with 100 μL of DMEM (2% FBS) containing the compound of interest at concentrations 50% greater than those indicated, including a DMSO control. Plates were then transferred into the BSL3 facility and 100 PFU (MOI = 0.025) was added in 50 μL of DMEM (2% FBS), bringing the final compound concentration to those indicated. Plates were then incubated for 48 hours at 37°C. After infection, supernatants were removed and cells were fixed with 4% formaldehyde for 24 hours prior to being removed from the BSL3 facility. The cells were then immunostained for the viral NP protein (an in-house mAb 1C7, provided by Dr. Thomas Moran) with a DAPI counterstain. Infected cells (488 nm) and total cells (DAPI) were quantified using the Celigo (Nexcelcom) imaging cytometer. Infectivity was measured by the accumulation of viral NP protein in the nucleus of the Vero E6 cells (fluorescence accumulation). Percent infection was quantified as ((Infected cells/Total cells) - Background) *100 and the DMSO control was then set to 100% infection for analysis. Data was fit using nonlinear regression and IC_50_s for each experiment were determined using GraphPad Prism software. Cytotoxicity was also performed using the MTT assay (Roche), according to the manufacturer’s instructions. Cytotoxicity was performed in uninfected VeroE6 cells with same compound dilutions and concurrent with viral replication assay. All assays were performed in biologically independent triplicates. The Vero E6 cell line used in this study is a kidney cell line; therefore, we cannot exclude that lung cells yield different results for some inhibitors (see also ‘Cells for RT-qPCR screening’ and ‘Antiviral drug and cell viability assays at Institut Pasteur’ for studies carried out at Institut Pasteur).

#### Phospholipidosis quantification in uninfected cells

Phospholipidosis was assessed as previously described (*24*). Briefly, A549 cells were cultivated in Ham’s F12-K Medium containing 10% FCS and seeded in a black 96-well plate with clear bottom at a density of 15000 cells per well. The day after seeding, the cells were treated for 24 hours with a dose-range of different drugs in presence of 7.5 µM NBD-PE (ThermoFisher, ref. N360). The final DMSO concentration was 0.2%. Amiodarone was used as a positive control for phospholipidosis. Before imaging, the cells were stained for 20 minutes at 37°C, 5% CO2 with a solution containing Hoechst (ThermoFisher, ref. H3570) (10 µg/ml) and Ethidium homodimer-2 (EthD-2; ThermoFisher, E3599) (2 µM) in complete culture medium for visualizing the total and dead cell populations respectively. Cells were washed once with pre-warmed HBSS +/+ and images were taken on an Arrayscan XTI (ThermoFisher) equipped with a 20x objective and the LED/filter combinations BGRFR_386_23, BGRFR_485_20 and BGRFR_549_15 for acquisition of Hoechst, NBD-PE and EthD-2 dyes respectively. The images were analyzed using the HCS Studio software (ThermoFisher). Briefly, the nuclei were detected using the Hoechst dye and the dead cell nuclei showing a co-staining with EthD-2 were excluded from the analysis. Then, the dots of NBD-PE were detected in the cytoplasm of each cell corresponding to a dilation of the nuclei to a maximum of 50 µm width and the total intensity of NBD-PE was measured in each cell for quantification of DIPL. After imaging, cytotoxicity was assessed doing an ATP quantification with the CellTiter-Glo® 2.0 Cell Viability assay kit (Promega, ref. G9241) following manufacturer’s instructions.

#### NBD-PE and Spike protein staining in infected cells

15000 A549-ACE2 cells per well were seeded into 96-well plates in DMEM (10% FBS) and incubated for 24 hours at 37°C, 5% CO2. Two hours prior to infection, the media was replaced with 100 μL of DMEM (2% FBS) containing the compound of interest at the indicated concentration and 7.5 µM NBD-PE dye (Invitrogen). At the time of infection, the media was replaced with virus inoculum (MOI 2 PFU/cell) and incubated for one hour at 37°C, 5% CO2. Following the adsorption period, the inoculum was removed, replaced with 100 μL of drug and NBD-PE-containing media, and cells incubated for an additional 24 hours at 37°C, 5% CO2. At this point, the supernatant was replaced with DMEM containing 10 µg/mL of Hoechst and 2 µM Ethidium homodimer-2 (Invitrogen) and incubated for 20 min at 37°C, 5% CO2. The cells were then washed once with HBSS+/+ and fixed with 4% (v/v) formalin in PBS. The plates were then imaged using the Opera Phenix High Content Screening System, taking 12 images per well with a 20x objective. Subsequently, the cells were washed and stained for Spike antigen, to identify infected cells. Briefly, the cells were permeabilized with Triton 0.1% for 10 minutes at room temperature and nonspecific staining was blocked with PBS 5% Bovine Serum Albumin (BSA) for two hours at room temperature. The cells were then stained with 1ug/mL of the anti-Spike human monoclonal antibody #48 (kindly provided by Hugo Mouquet, Institut Pasteur-Paris) overnight at 4°C and with a goat anti-human secondary antibody AlexaFluor647 for one hour at room temperature. Upon staining, the cells were imaged once more in the same fields of view using the Opera Phenix screening system.

For image analysis the Columbus image analysis (PerkinElmer) system was used. Nuclei touching the border of the image were rejected. Living cells (not stained with Ethidium homodimer 2) were identified and the total intensity of NBD-PE dots in a 50 µm radius circle centered on the nucleus of living cells was measured, using the spot detection algorithm. To quantify Sars-CoV-2 infection, the area of Spike+ staining was quantified and normalized by the number of nuclei using the same software. Two-way ANOVAs were performed on pooled data from three biological replicates using GraphPad Prism.

#### General Chemical Synthesis Procedure

Anhydrous solvents were purchased from Acros Organics. Unless specified below, all chemical reagents were purchased from Sigma-Aldrich and AK Scientific. Commercial solvents and reagents were used as received. All reactions were performed in oven-dried glassware fitted with rubber septa under a positive pressure of argon, unless otherwise noted. Air- and moisture-sensitive liquids were transferred via syringe. Solutions were concentrated by rotary evaporation at or below 40 °C. Analytical thin-layer chromatography (TLC) was performed using glass plates pre-coated with silica gel (0.25-mm, 60-Å pore size, 230−400 mesh, Merck KGA) impregnated with a fluorescent indicator (254 nm). TLC plates were visualized by exposure to ultraviolet light (UV), then were stained by submersion in a 10% solution of phosphomolybdic acid (PMA) in ethanol or an acidic ethanolic solution of p-anisaldehyde followed by brief heating on a hot plate. This solution was prepared by sequential additions of concentrated sulfuric acid (5.0 mL), glacial acetic acid (1.5 mL) and p-anisaldehyde (3.7 mL) to absolute ethanol (135 mL) at 23 °C with efficient stirring. Flash column chromatography was performed with Teledyne ISCO CombiFlash EZ Prep chromatography system, employing pre-packed silica gel cartridges (Teledyne ISCO RediSep).

#### Instrumentation

Proton nuclear magnetic resonance (^1^H NMR) spectra and carbon nuclear magnetic resonance (^13^C NMR) spectra were recorded on Bruker AvanceIII HD 2-channel instrument (400 MHz/100 MHz) at 23 °C. Proton chemical shifts are expressed in parts per million (ppm, δ scale) and are referenced to residual protium in the NMR solvent (CHCl_3_: δ 7.26, D_2_HSOCD_3_: δ 2.50). Carbon chemical shifts are expressed in parts per million (ppm, δ scale) and are referenced to the carbon resonance of the NMR solvent (CDCl_3_: δ 77.0, CD_3_SOCD_3_: δ 39.5). Data are represented as follows: chemical shift, multiplicity (s = singlet, d = doublet, t = triplet, q = quartet, dd = doublet of doublets, dt = doublet of triplets, m = multiplet, br = broad, app = apparent), integration, and coupling constant (J) in Hertz (Hz). High-resolution mass spectra were obtained using a Waters Xevo G2-XS time-of-flight mass spectrometer. Unless otherwise specified, diastereomeric ratios of products are reported as (major diastereomer) : (sum of minor diastereomers).

#### ZZY-10-051

N-Boc-4-piperidone (36.8 mg, 0.185 mmol) and sodium cyanoborohydride (11.6 mg, 0.185 mmol) were added sequentially to a stirred suspension of 1-[3-(5-methoxytetralin-1-yl)propyl]piperazine hydrochloride (S1•HCl, 20.0 mg, 0.0616 mmol) in methanol (0.50 mL). The mixture was stirred at 23 °C and the reaction progress was monitored by LC-MS. After 24 h, the reaction mixture was partitioned between saturated aqueous sodium bicarbonate solution (5 mL) and dichloromethane (5 mL). The layers were separated, and the aqueous layer was extracted with dichloromethane (2 x 5 mL). The combined organic layers were dried over sodium sulfate. The dried solution was filtered, and the filtrate was concentrated. The residue was purified by column chromatography (0–10% methanol–dichloromethane, 4-g RediSep(R) Rf column, Teledyne ISCO, Lincoln, NE) to afford the product as a white foam (26.0 mg, 90%). HRMS (ESI^+^): Calculated for [C_28_H_45_N_3_O_3_+H]^+^: 472.3539. Found: 472.3518.

#### Correlation analyses

Compound-by-compound phospholipidosis versus SARS-CoV-2 infection correlation analyses were performed using the “correlation” function in GraphPad Prism using raw RT-qPCR values normalized to DMSO from the same experiment at 100%. To model the pooled correlation we used Bayesian inference through the brms R package v2.14.4 (*67*) the log PLD. With weakly informative priors, fitting 107 drug/concentration pairs yielded posterior parameter means and 95% credible interval (95%CI) estimates of IC_50_: 43 [38, 48]%, hill: -5.6 [-7.0, -4.5], and Sigma 2.0 [0.14, 1.78]. Bayesian leave-one-out *R^2^* values were computed by using the loo package v2.4.1(*68*).

#### Literature search for SARS-CoV-2 repurposed antivirals

Twelve major phenotypic antiviral repurposing papers were sourced from the literature(*4, 15–17, 26, 28–34*). Drug names were selected from each paper based on the author-reported number of *in vitro* hits at the most strict reported threshold. If an explicit mention of the number of hits was not mentioned, all molecules with demonstrated dose-response inhibition of SARS-CoV-2 were selected. SMILES for each compound were retrieved from PubChem(*69*) using the PubChemPy API (https://pubchempy.readthedocs.io). Using the PubChem canonical SMILES, each molecule’s cLogP was calculated using RDkit-2019.09.3.0 (http://www.rdkit.org), and JChem’s cxcalc command line tool was used to calculate each molecule’s most basic pKa, JChem-15.11.23.0, ChemAxon (https://www.chemaxon.com). Molecules with cLogP ≥ 3 and pKa ≥ 7.4 were considered CADs. ECFP-4-based Tanimoto coefficients (Tcs) were calculated for the list of CADs to all molecules from a list of known phospholipidosis inducers (*19, 25*), and the maximum Tc for each CAD to a known phospholipidosis inducer was used for filtering the CAD list for known, and highly likely, phospholipidosis inducers (Tc ≥ 0.4).

#### Pharmacokinetics

Pharmacokinetic experiments were performed by Bienta (Enamine Biology Services). Plasma pharmacokinetics and lung distribution for amiodarone, sertraline, PB28, tamoxifen, and elacridar were investigated following five intraperitoneal (i.p.) doses of each drug in 21 male CD-1 mice per drug condition plus 1 mouse per vehicle condition (106 mice per drug experiment total). The formulations for each compound was as follows: amiodarone-PG-PEG400 (80%:20%); sertraline-DMA-PEG400-physiological saline (20%:20%:60%); PB28-PG-PEG400 (80%:20%); tamoxifen-corn oil (100%); Elacridar-DMA-PEG400-2HPβCD-water (25%:25%:25%:25%). Mice were injected i.p. with 2,2,2-tribromoethanol at the dose of 150 mg/kg prior to drawing blood. Blood collection was performed from the orbital sinus in microtainers containing K2EDTA at 0, 5, 15, 30, 60, 120, 360, and 1440 minutes after drug injection. Animals were sacrificed by cervical dislocation after the blood samples collection. After this, lung samples were collected and weighted. All samples were immediately processed, flash-frozen and stored at -70°C until subsequent analysis.

Plasma samples (50 μL) were mixed with 200 μL of internal standard (IS) solution. After mixing by pipetting and centrifuging for 4 min at 6000 rpm, 0.5 μL of each supernatant was injected into the LC-MS/MS system. Solution of each compound (400 ng/ml in acetonitrile-methanol mixture, 1:1, v/v) was used as IS for drug quantification in plasma samples. Lung samples were dispersed in 3.5 volumes of IS400(90) using stainless steel beads (115 mg ± 5 mg) in The Bullet Blender® homogenizer for 90 sec at speed 10. After this, the samples were centrifuged for 4 min at 14,000 rpm, and 0.5 μL of each supernatant was injected into the LC-MS/MS system. Solution of a reference compound (sertraline or amiodarone; 400 ng/ml in water-methanol mixture, 1:9, v/v) was used as IS for quantification of the drugs in lung samples.

Analyses of plasma and lung samples were conducted at Enamine/Bienta. The concentrations of drugs in plasma and lung samples were determined using high performance liquid chromatography/tandem mass spectrometry (HPLC-MS/MS). The Shimadzu HPLC system used consists of two isocratic pumps LC-10ADvp, an autosampler SIL-30AC MP, a sub-controller FCV-20AHs and a degasser DGU-14A. Mass spectrometric analysis was performed using a 4000 Q TRAP instrument from MDS Sciex (Canada) with an electro-spray (ESI) interface. The data acquisition and system control was performed using Analyst 1.6.3 software from AB Sciex. The concentrations of the test compound below the lower limit of quantitation (LLOQ, amiodarone: 5 ng/mL for plasma and 7 ng/g for lung; sertraline: 10 ng/mL for plasma and 17.5 ng/g for lung; PB28: 5 ng/mL for plasma and 20 ng/g for lung; tamoxifen: 10 ng/mL for plasma and 8 ng/g for lung; elacridar: 5 ng/mL for plasma and 35 ng/g for lung) were designated as zero. The pharmacokinetic data analysis was performed using noncompartmental, bolus injection or extravascular input analysis models in WinNonlin 5.2 (PharSight). Data below LLOQ were presented as missing to improve validity of T½ calculations.

#### Animal models of SARS-CoV-2 infection experiments

All the antiviral studies were performed in an animal biosafety level 3 (BSL3) facility at the Icahn school of Medicine in Mount Sinai Hospital, New York City. All work was conducted under protocols approved by the Institutional Animal Care and Use Committee (IACUC). We used a model of BALB/c mice transduced intranasally with an adenovirus expressing human ACE2 (hACE2) for all experiments. We used 29 female 12-week-old specific pathogen–free BALB/c mice (the Jackson laboratory strain 000651). Five days prior to infection with SARS-CoV-2, BALB/c mice were infected intranasally with 2.5x10^8^ PFU of an adenovirus carrying the gene for hACE2. Viral seed stocks for non-replicating E1/E3 deleted viral vectors based on human adenovirus type-5 expressing the human angiotensin-converting enzyme 2 (Ad-ACE2) receptor under the control of a CMV promoter, were obtained from the Iowa Viral Vector Core Facility. Viral stocks were amplified to high titers following infection of T-Rex TM-293 cells and purification using two sequential rounds of cesium chloride (CsCl) ultracentrifugation, as described previously (*70*), (*71*). The infectious titer was determined using a tissue culture infectious dose-50 (TCID_50_) end-point dilution assay, and physical particle titer quantified by micro-bicinchoninic acid (microBCA) protein assay, both described previously (*70*). Remdesivir was administered subcutaneously (s.c.) while the other five remaining drugs were injected intraperitoneally (i.p.), with amiodarone and elacridar dosed once per day and remdesivir, PB28, tamoxifen, and sertraline being dosed twice per day, for a total of 3 days, consistent with their pharmacokinetic profiles. We administered vehicle (PBS, i.p., twice-daily) or drug treatments, two hours before intranasal infection with 1 × 10^4^ PFU of SARS-CoV-2 in 50 μL of PBS. Mice were anesthetized with a mixture of ketamine/xylazine before each intranasal infection. Each day mouse body weight was measured. Three days post infection (dpi) animals were humanely euthanized. Whole left lungs were harvested and homogenized in PBS with silica glass beads then frozen at −80°C for viral titration via TCID_50_. Briefly, infectious supernatants were collected at 48h post infection and frozen at −80°C until later use. Infectious titers were quantified by limiting dilution titration using Vero E6 cells. Briefly, Vero E6 cells were seeded in 96-well plates at 20,000 cells/well. The next day, SARS-CoV-2-containing supernatant was applied at serial 10-fold dilutions ranging from 10 −1 to 10 −6 and, after 5 days, viral cytopathic effect (CPE) was detected by staining cell monolayers with crystal violet. TCID_50_/mL were calculated using the method of Reed and Muench. The Prism software (GraphPad) was used to determine differences in lung titers using an unpaired T test.

#### COVID-19 clinical trial expenditure analysis

Data for COVID-19 clinical trials were downloaded from the DrugBank(*56*) COVID-19 dashboard (accessed on 2021-02-16). Supplemental information for each compound, including its SMILES and DrugBank ID was downloaded and unpacked from the DrugBank “All drugs” 2021-01-03 release(*72*). A total of 3395 treatments were recorded in the COVID-19 dashboard at the time of download, and 2244 of those annotations were for small molecules from a total of 1490 unique clinical trials worldwide. Using the DrugBank annotated SMILES, each molecule’s cLogP was calculated using RDkit-2019.09.3.0 (http://www.rdkit.org), and JChem’s cxcalc command line tool was used to calculate each molecule’s most basic pKa, JChem-15.11.23.0, ChemAxon (https://www.chemaxon.com). Molecules with cLogP ≥ 3 and pKa ≥ 7.4 were considered CADs. Azithromycin, a non-CAD phospholipidosis inducer, was also included in the CAD dataset due to its shared mechanism of action with CAD antivirals whereas fluvoxamine, a CAD known to act through immune-mediated processes was excluded from the CAD dataset. Additional subsets of these data, including filtering out chloroquine and hydroxychloroquine trials from the CADs, were also analyzed. To quantify the estimated cost of clinical trials, the molecules from a given subset were first filtered for unique clinical trial IDs, then were grouped based on clinical trial phase and multiplied by the average cost of an anti-infective clinical trial in that phase(*60, 61*). Mixed-phase trials were multiplied by the cost of the more advanced phase only.

---

**Fig. S1.**
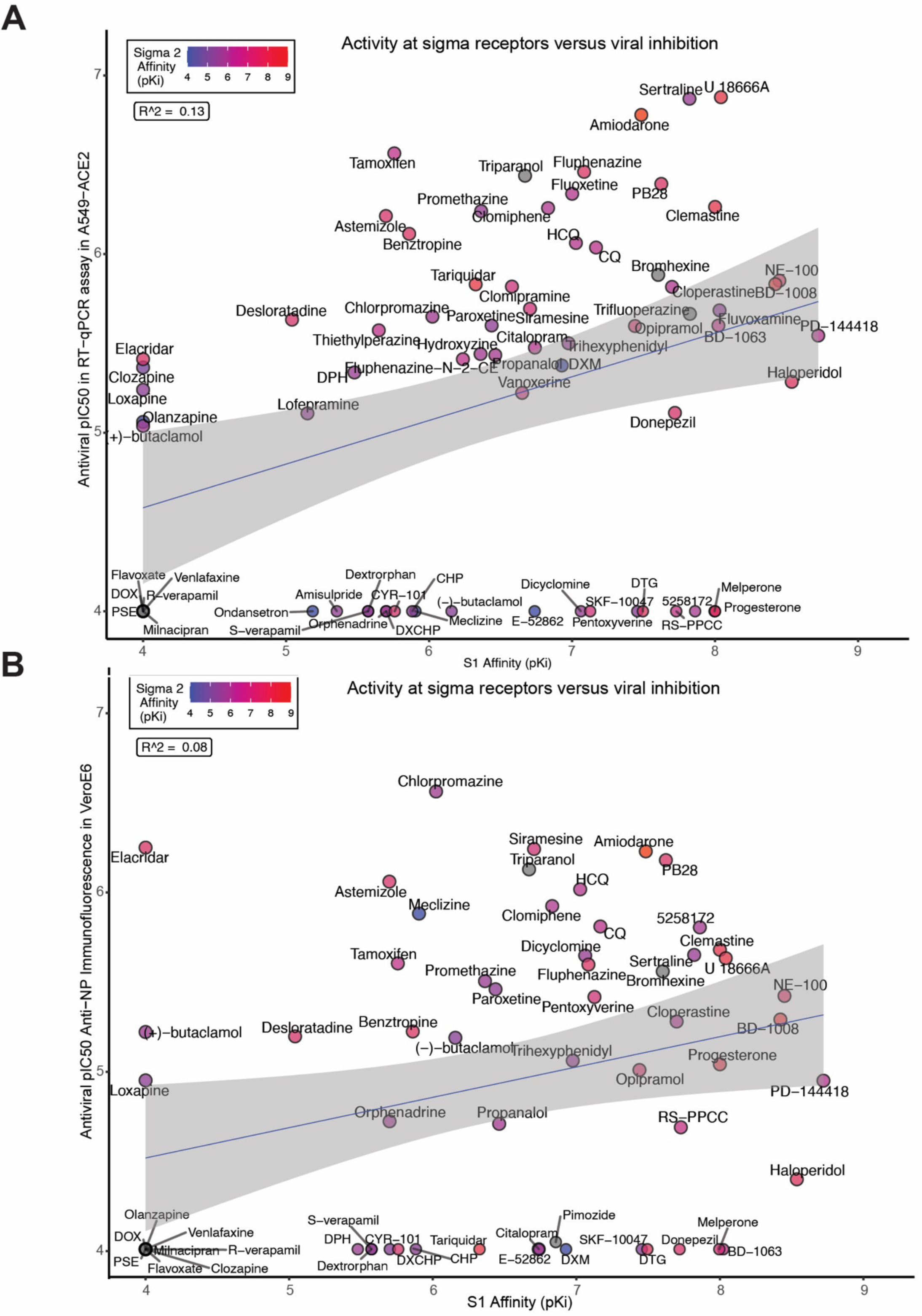
Correlation analyses for sigma receptor affinity and antiviral activity. **A.** pKi at sigma-1 was correlated with pIC50 in the RT-qPCR assay in A549-ACE2 cells. pKi at sigma-2 is denoted by the colors. **B**. pKi at sigma-1 was correlated with pIC50 in the anti-NP immunofluorescence viral infectivity assay in VeroE6 cells. Abbreviations: CQ: chloroquine; HCQ: hydroxychloroquine; DOX: doxylamine; PSE: pseudephedrine; CHP: chlorpheniramine; DXCHP: dexchlorpheniramine; DXM: dextromethorphan; DPH: diphenhydramine; DTG: ditolylguanidine.

**Fig. S2.**
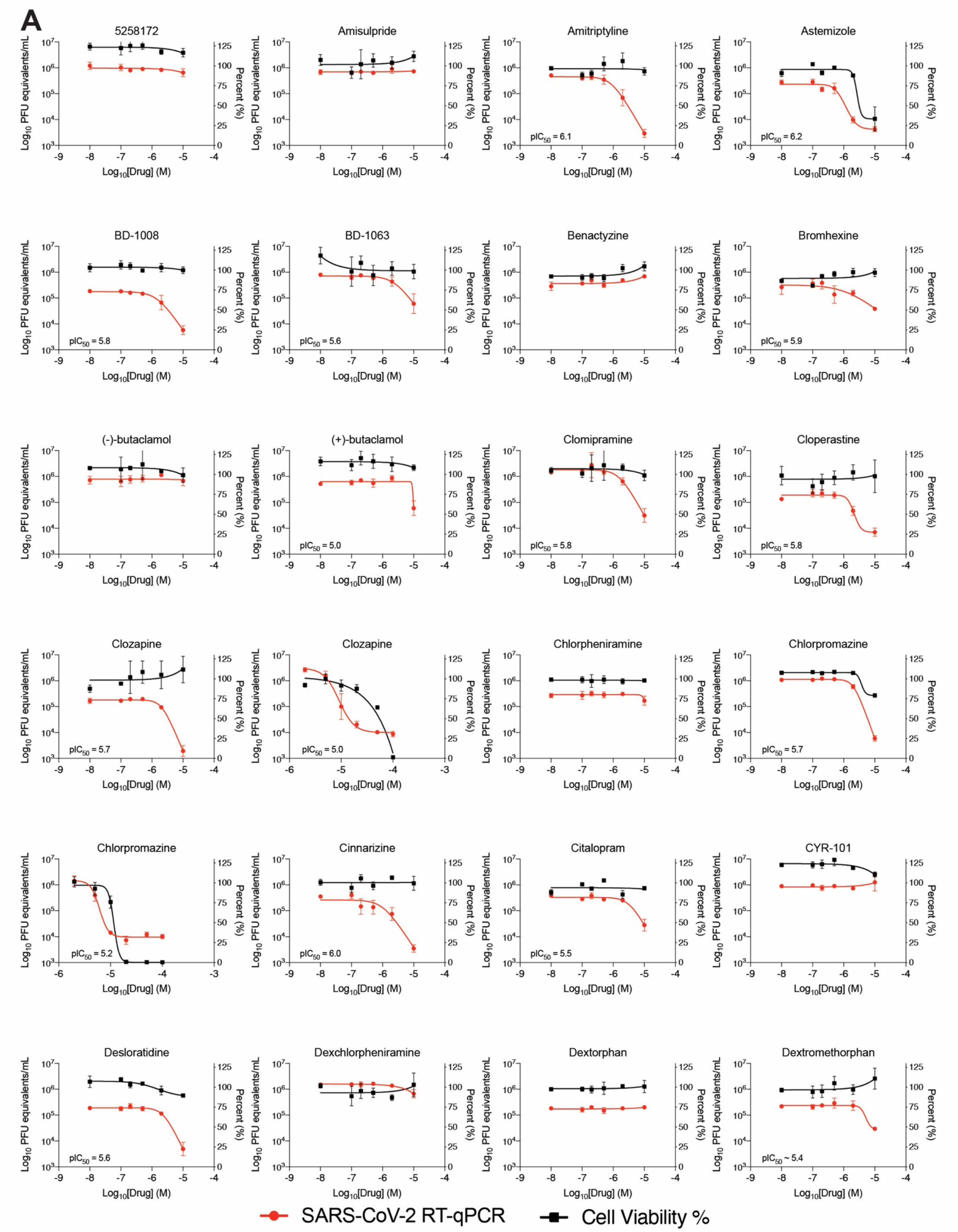
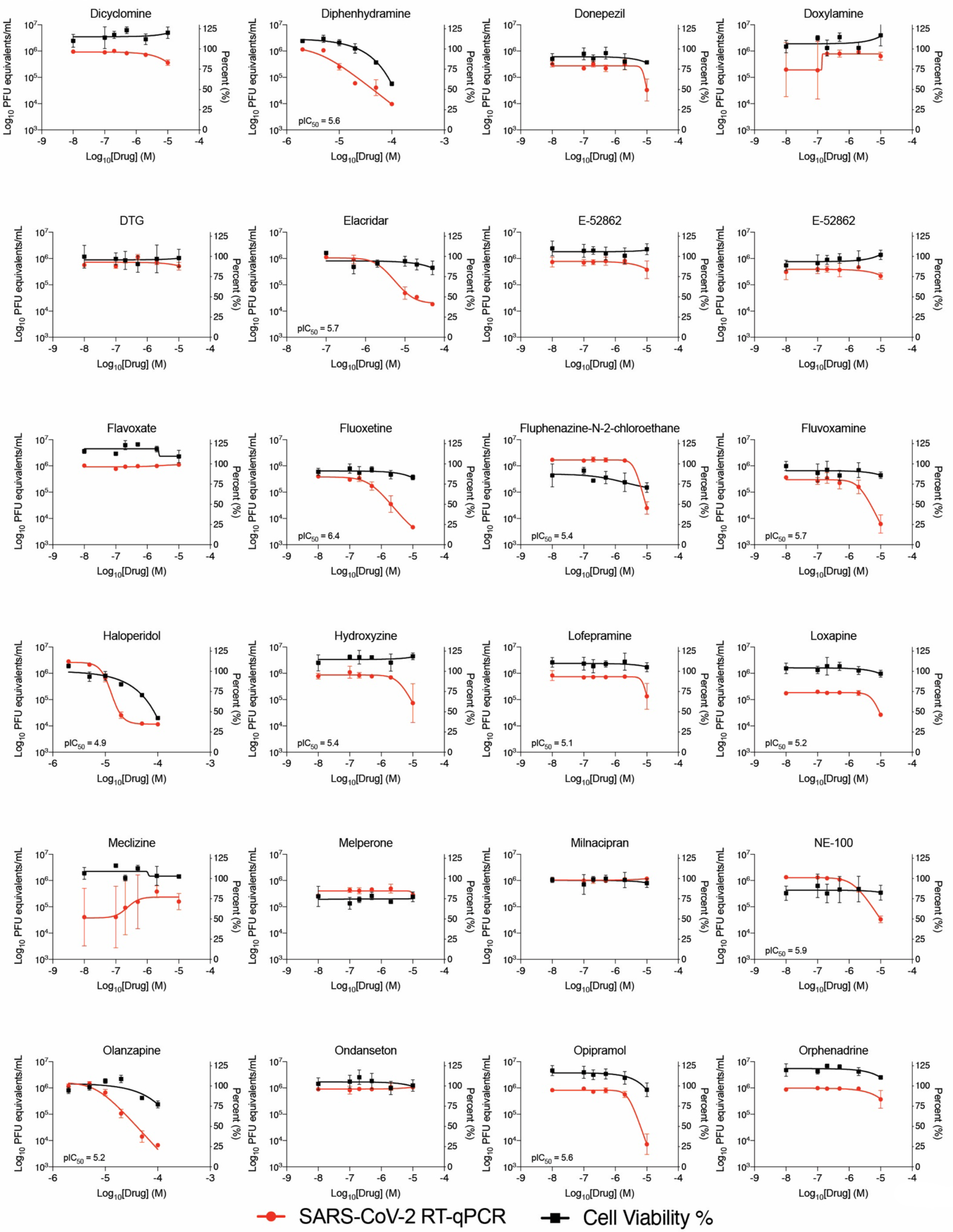
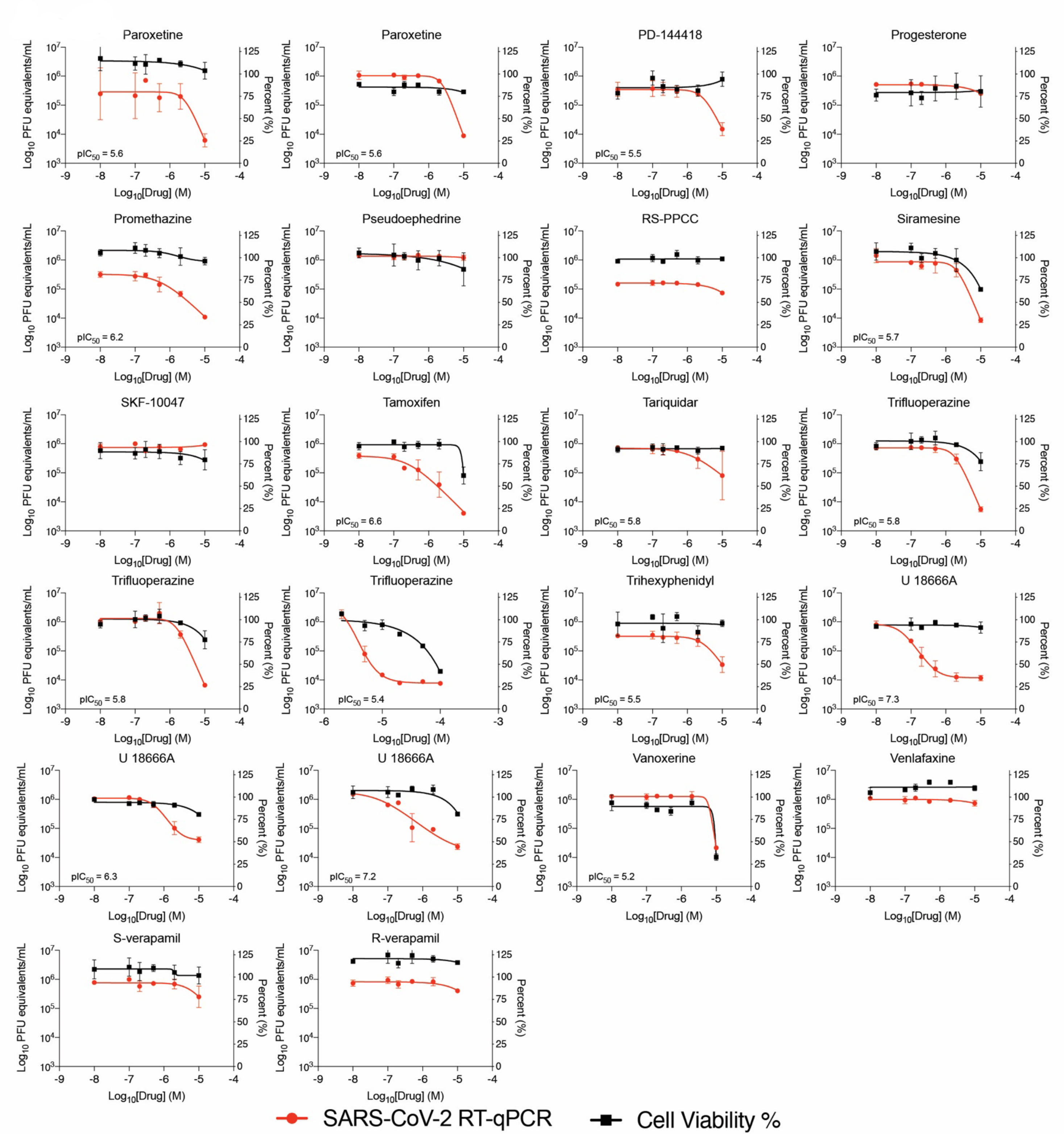
Dose response curves for a set of cationic amphiphilic drugs in an RT-qPCR viral infectivity assay. **A.** Viral infectivity and cell viability data for a subset of literature-identified cationic amphiphilic drugs in VeroE6 cells. Data shown are mean ± SD from three technical replicates. Biological replicates are shown as separate graphs when available.

**Fig. S3.**
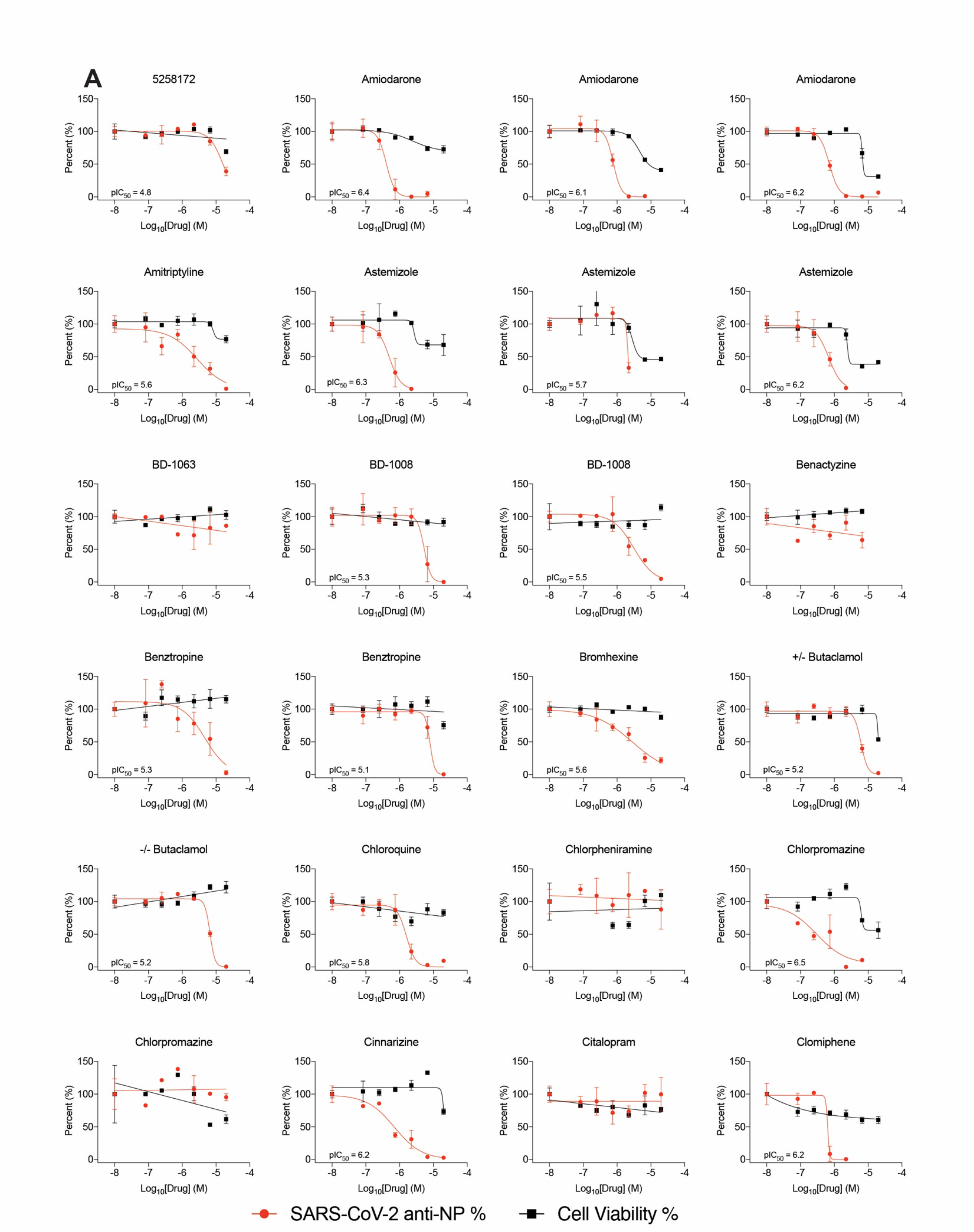
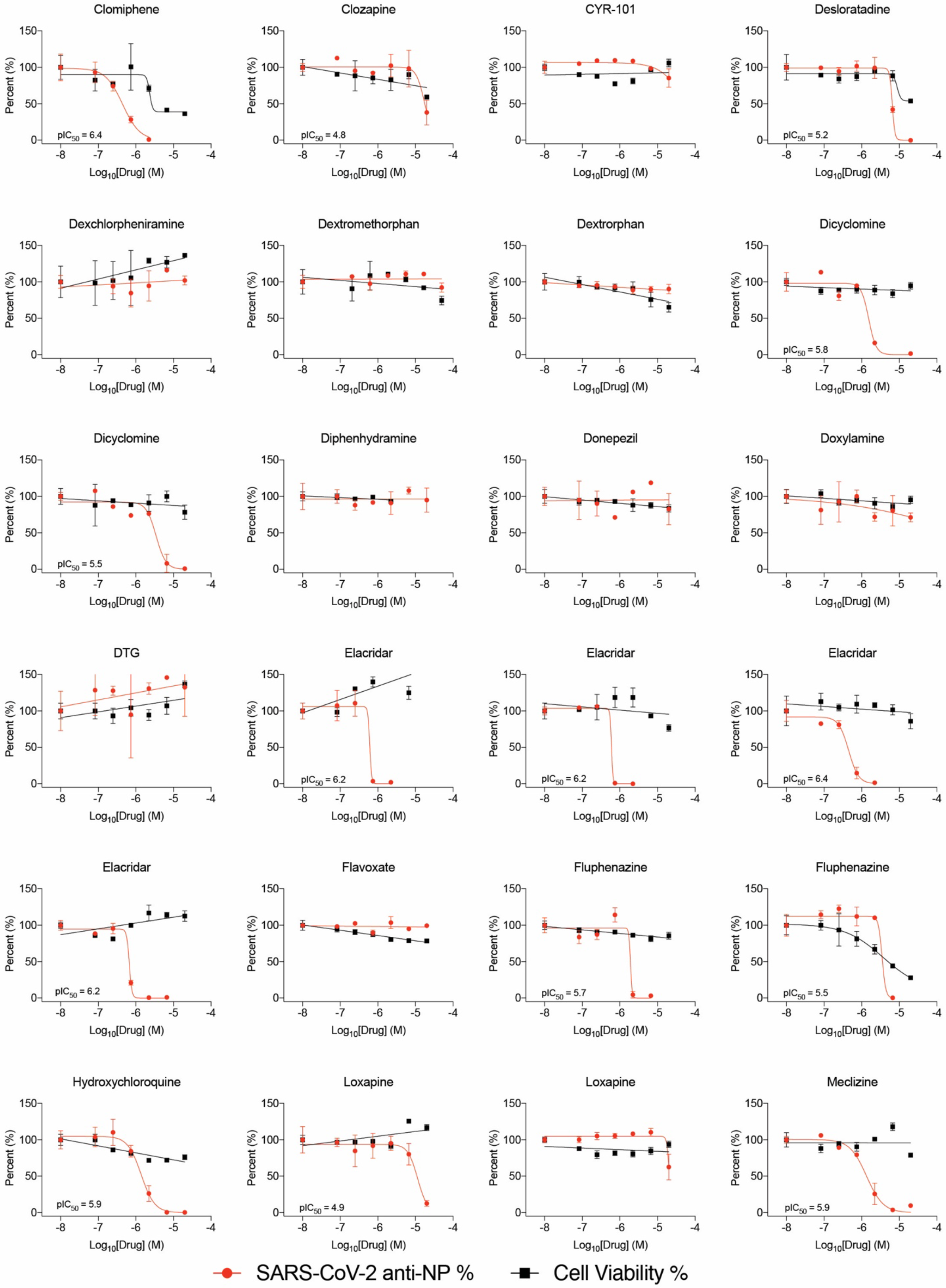
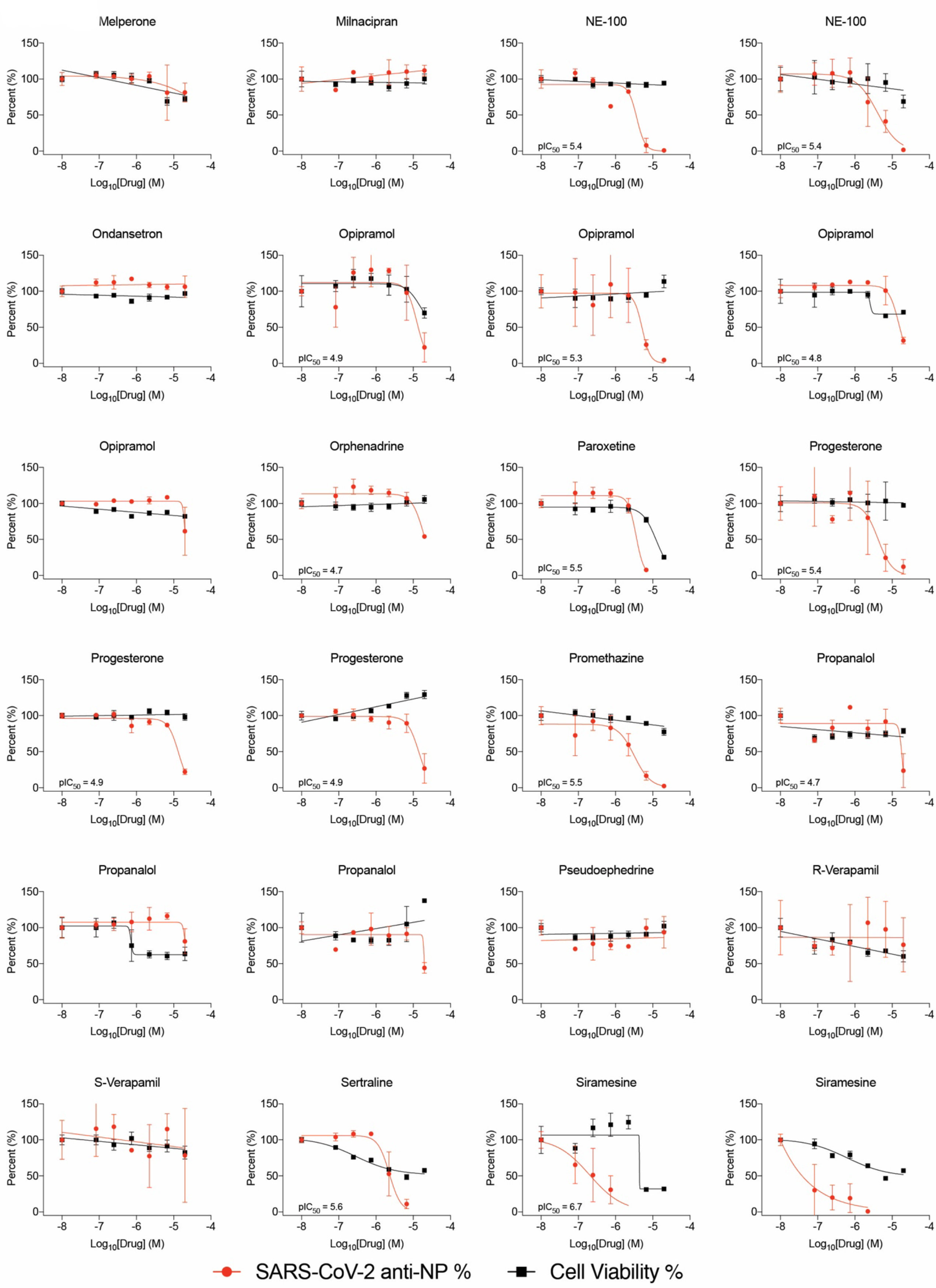
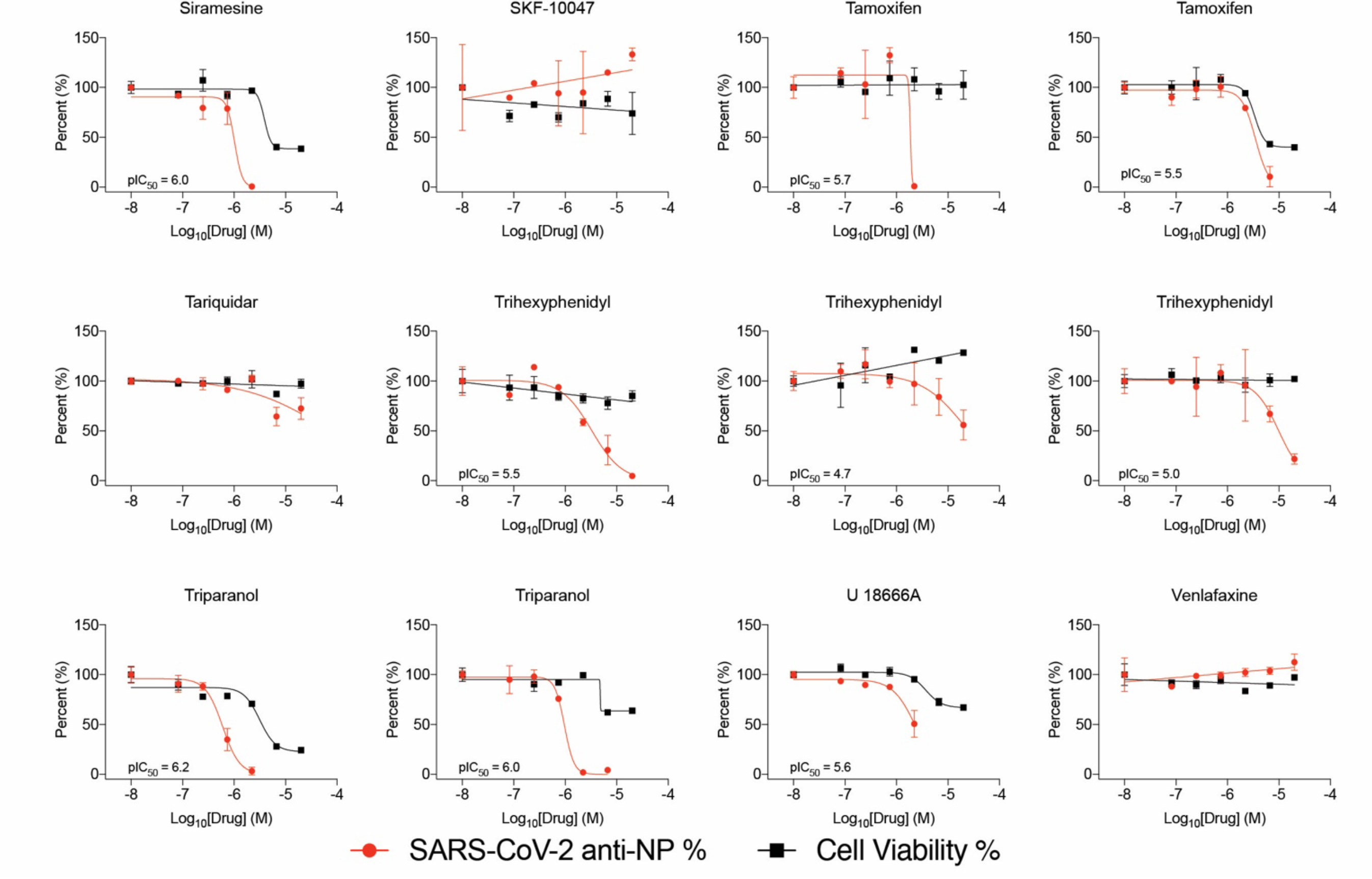
Dose response curves for a set of cationic amphiphilic drugs (CADs) in an anti-NP immunofluorescence viral infectivity assay. **A.** Viral infectivity and cell viability data for a subset of literature-identified CADs in VeroE6 cells. Data shown are mean ± SD from three technical replicates. Biological replicates are shown as separate graphs when available.

**Fig. S4.**
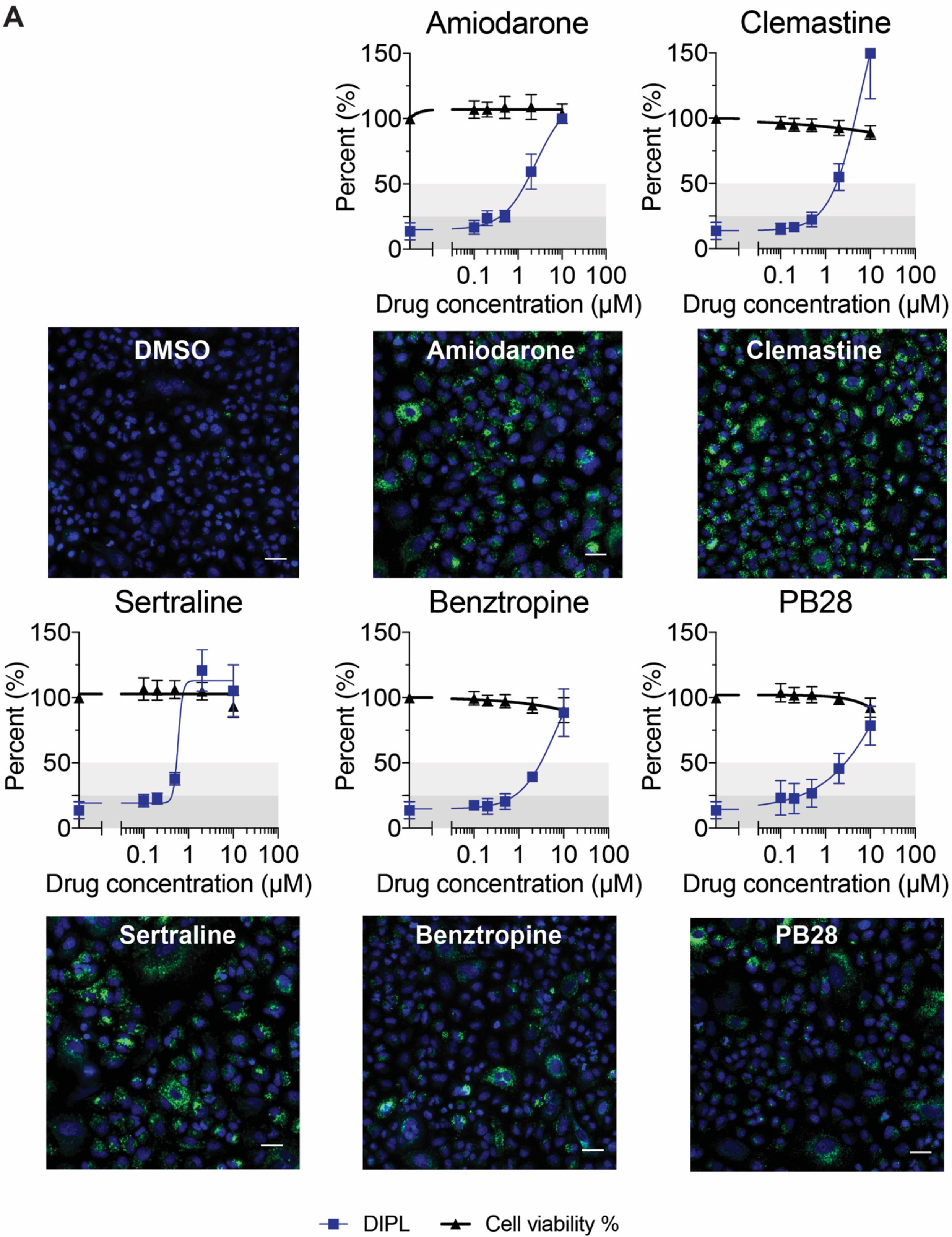
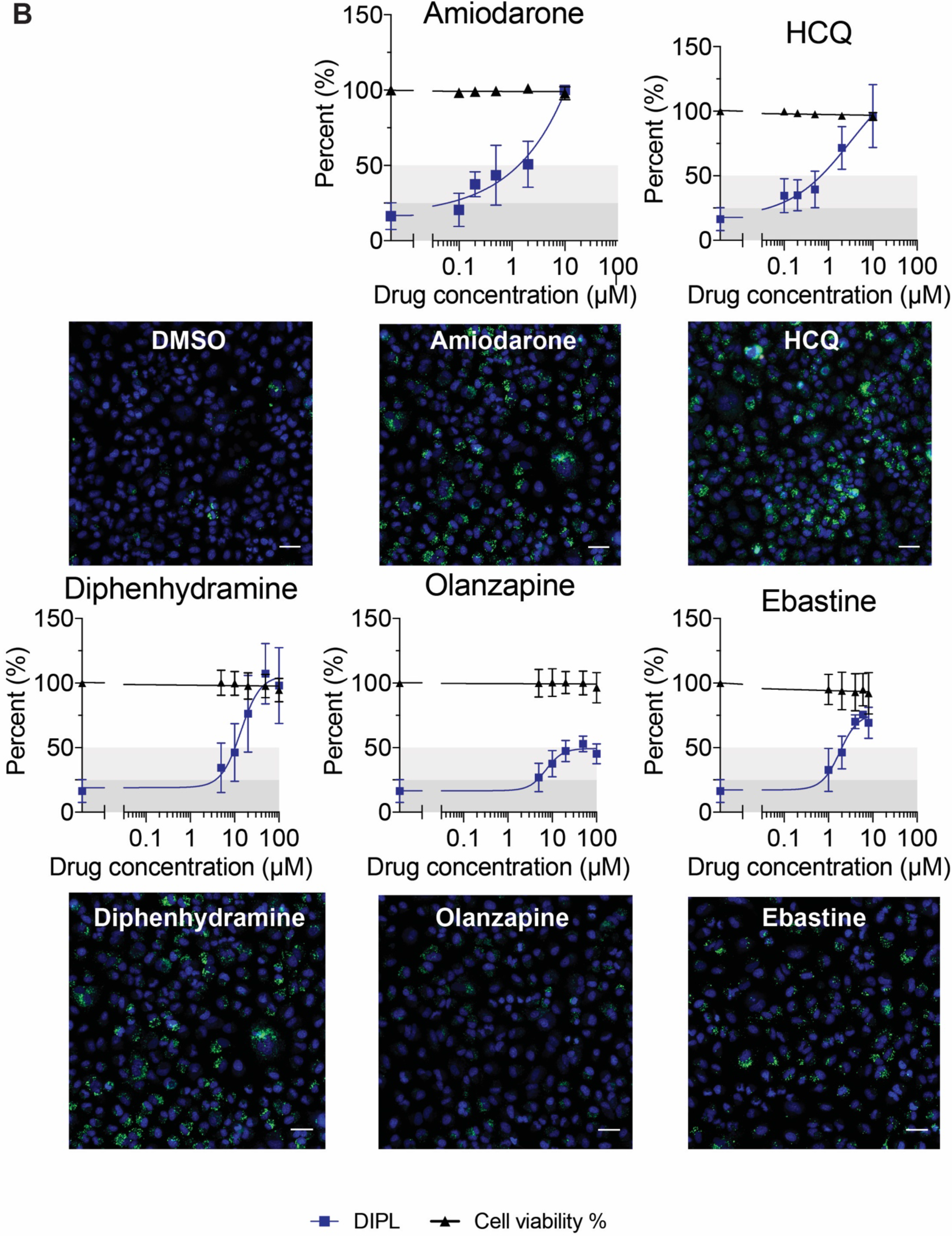
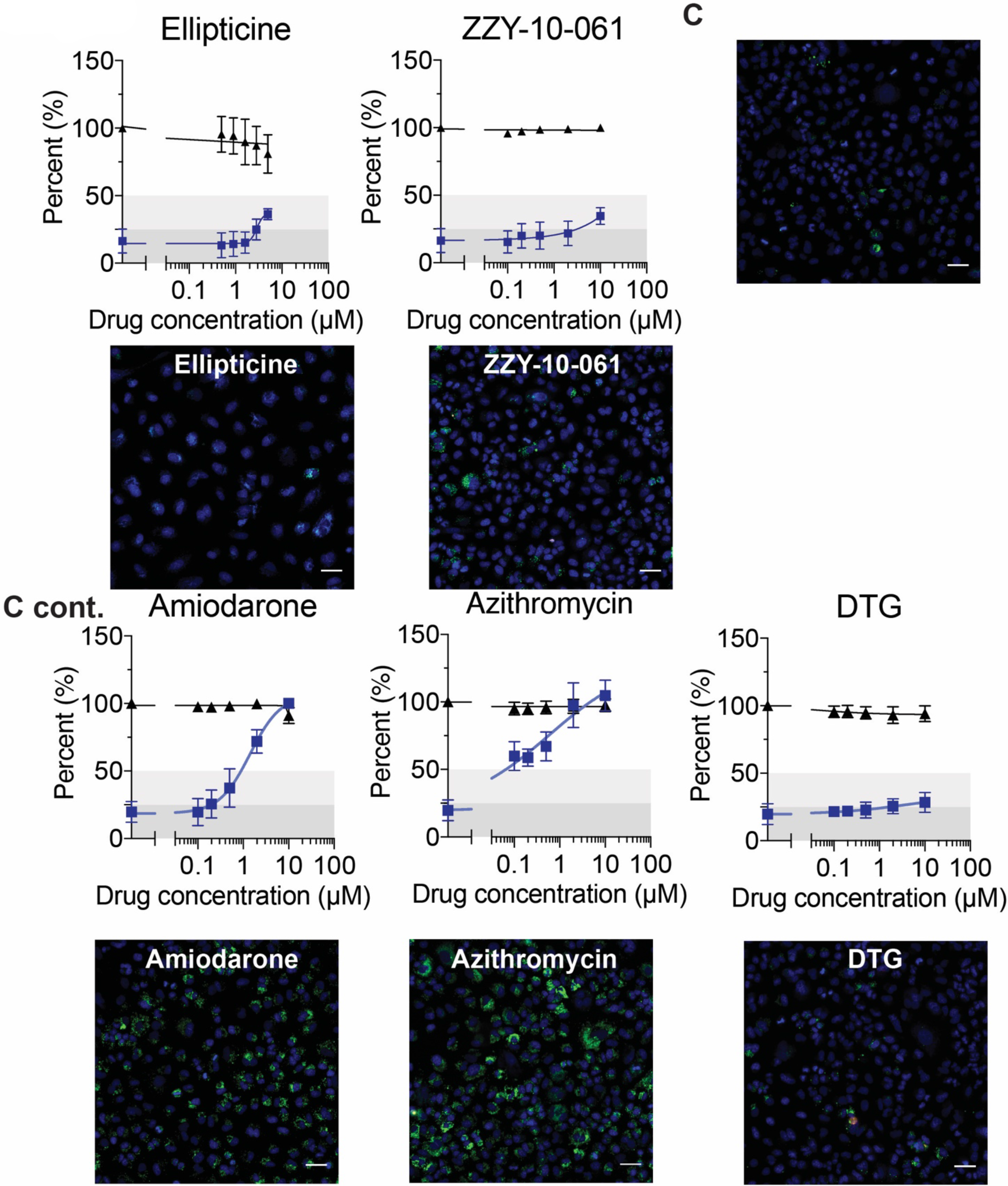
Dose response curves for drugs measured in the phospholipidosis and cell viability assays and plate images at top tested concentrations from batch 1- **A.**, batch 2- **B.,** and batch 3- **C.** Blue = Hoechst nuclei staining, Green = NBD-PE phospholipid staining, Red = EthD-2 staining for dead cells. Dose response of NBD-PE aggregation was normalized to DMSO and 10 µM amiodarone from the same batch, and cell viability was normalized to DMSO. Data shown are pooled means ± SD from three biological and three technical replicates.

**Fig. S5.**
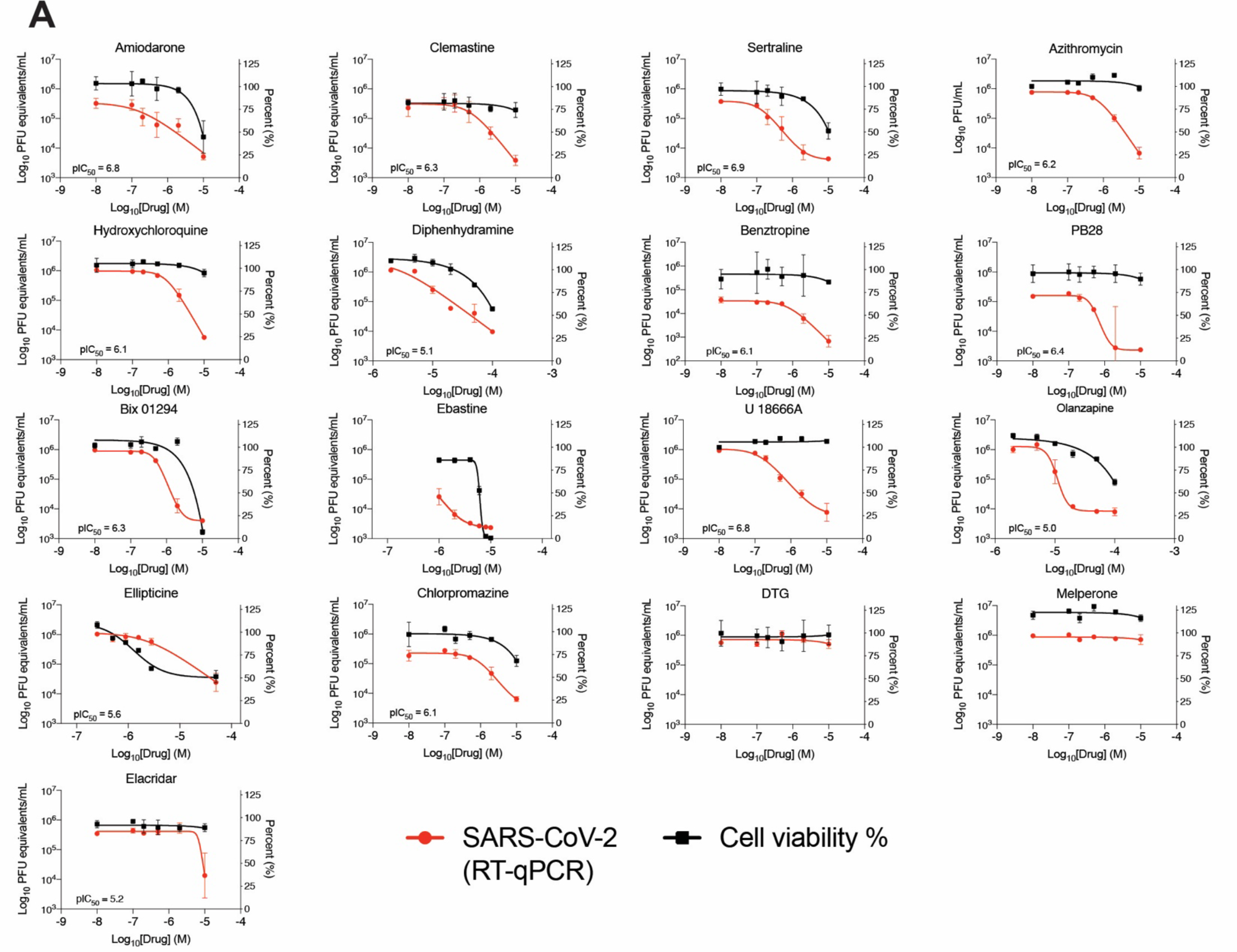
Dose response curves for cationic amphiphilic drugs in the RT-qPCR viral infectivity assay that were measured for NBD-PE aggregation. **A.** Viral infectivity and cell viability data for a subset of drugs that were selected for the DIPL correlation analysis. Data shown are mean ± SD from three technical replicates. The concentrations from these experiments match what was tested in the NBD-PE assay. Data for amiodarone, chlorpromazine, and hydroxychloroquine are reprinted from Gordon *et al.* with permission (*15*). Data for ZZY-10-051 and ZZY-10-061 are in **Fig. S7**.

**Fig S6.**
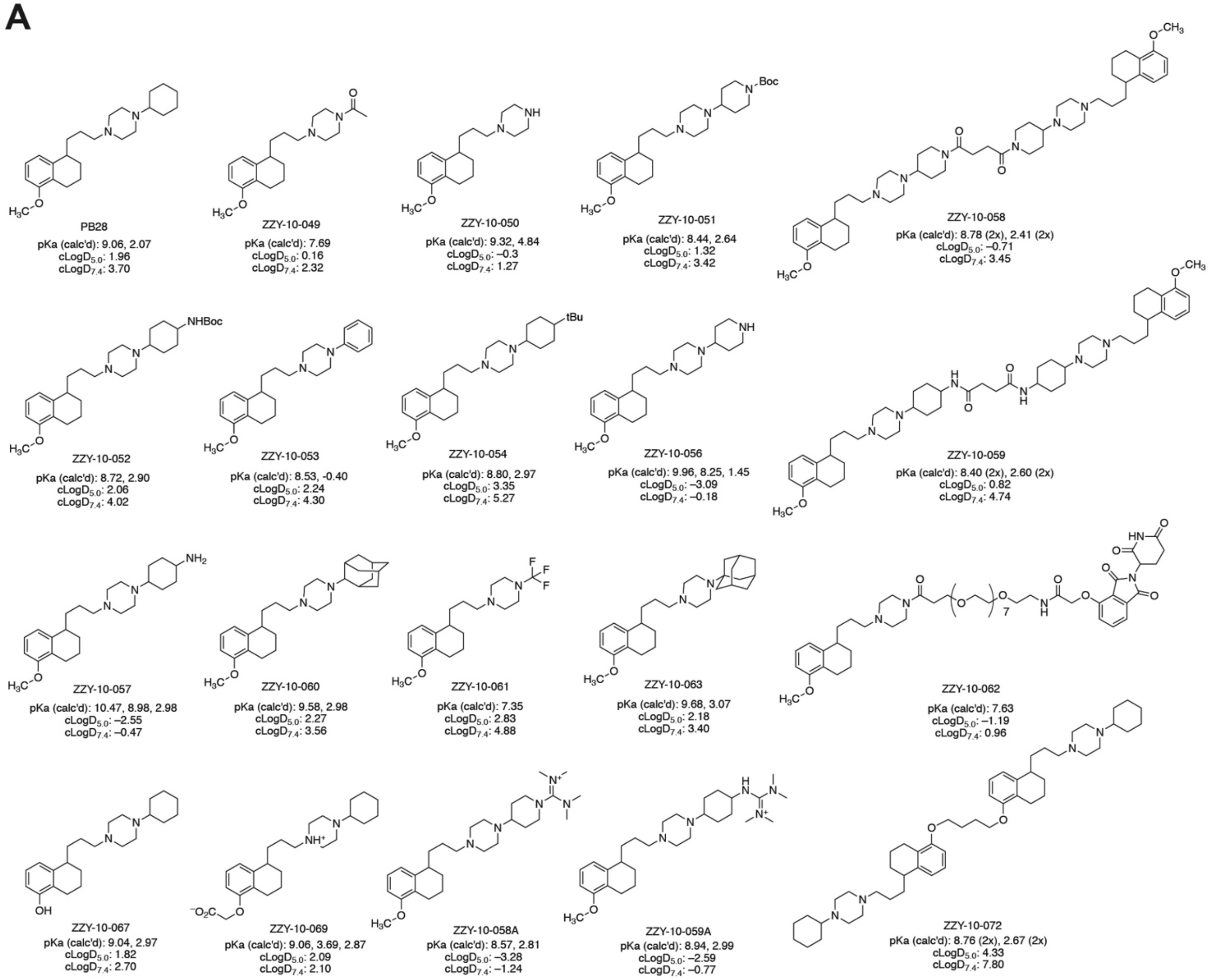
PB28 analog structures. **A.** Chemical structures of PB28 analogs tested in this study, shown in their neutral form. All compounds were prepared as racemates. Compounds ZZY-10-061, ZZY-10-062, ZZY-10-064, ZZY-10-056, ZZY-10-057, ZZY-10-058, ZZY-10-059 and ZZY-10-072 contain mixtures of diastereomers that were not resolved or separated. With the exception of ZZY-10-061 and ZZY-10-062, all compounds were prepared as HCl salts by acidification of their neutral forms with an ethereal solution of hydrogen chloride, or by lyophilization of their aqueous solutions in 50 mM HCl.

**Fig S7.**
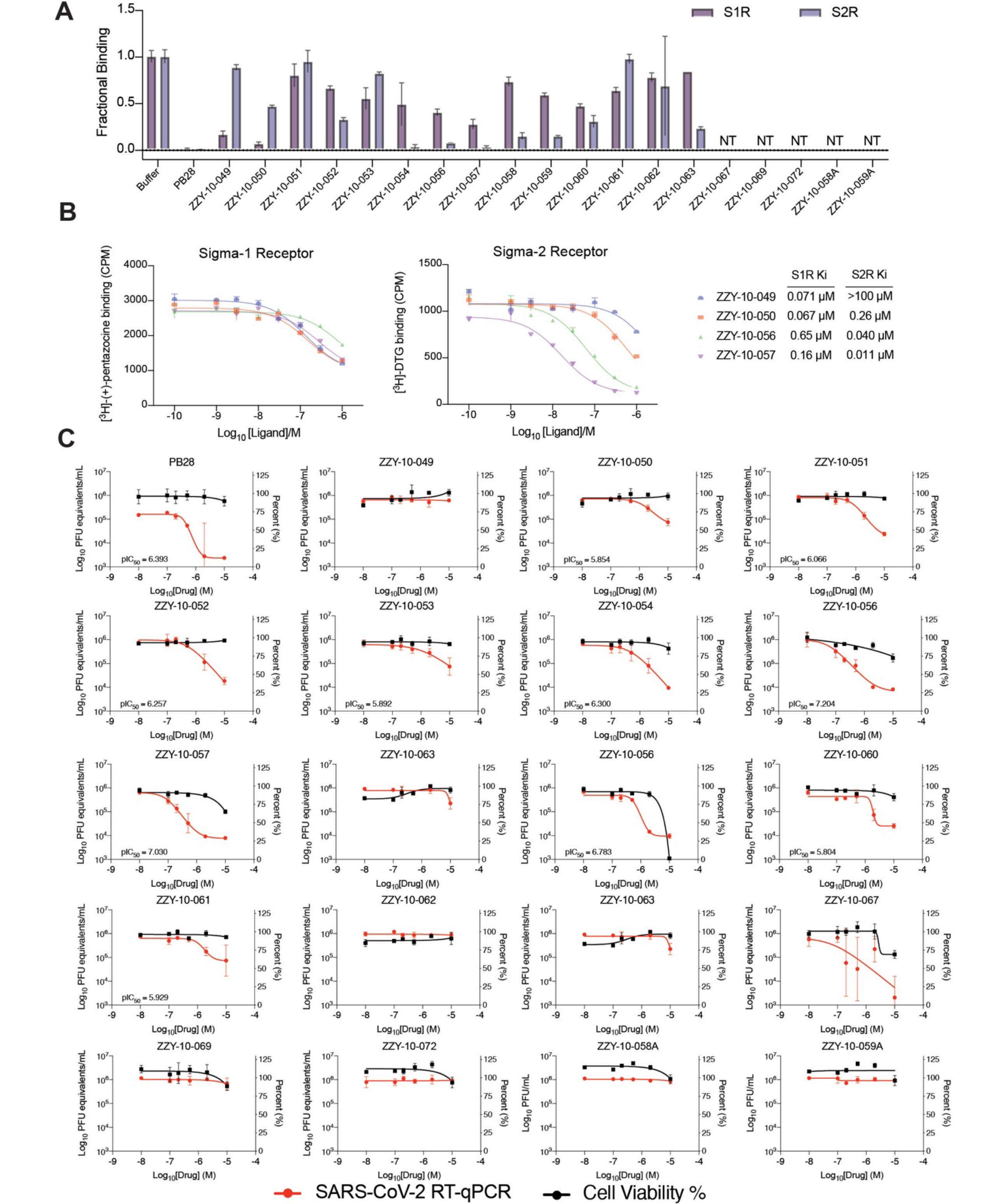
PB28 analog antiviral and sigma binding data. **A.** Fractional binding of PB28 analogs against sigma-1 (purple) and sigma-2 (blue) normalized to a buffer control at 1.0 in a radioligand binding experiment. Data shown are mean ± SEM from three technical replicates. **B.** Dose-response curves for selected PB28 analogs in sigma-1 and sigma-2 radioligand competition binding assay. Data points shown are mean ± SEM from three technical replicates. **C.** Viral infectivity data for PB28 analogs A549-ACE2 cells. Data shown are mean ± SD from three technical replicates.

**Fig S8.**
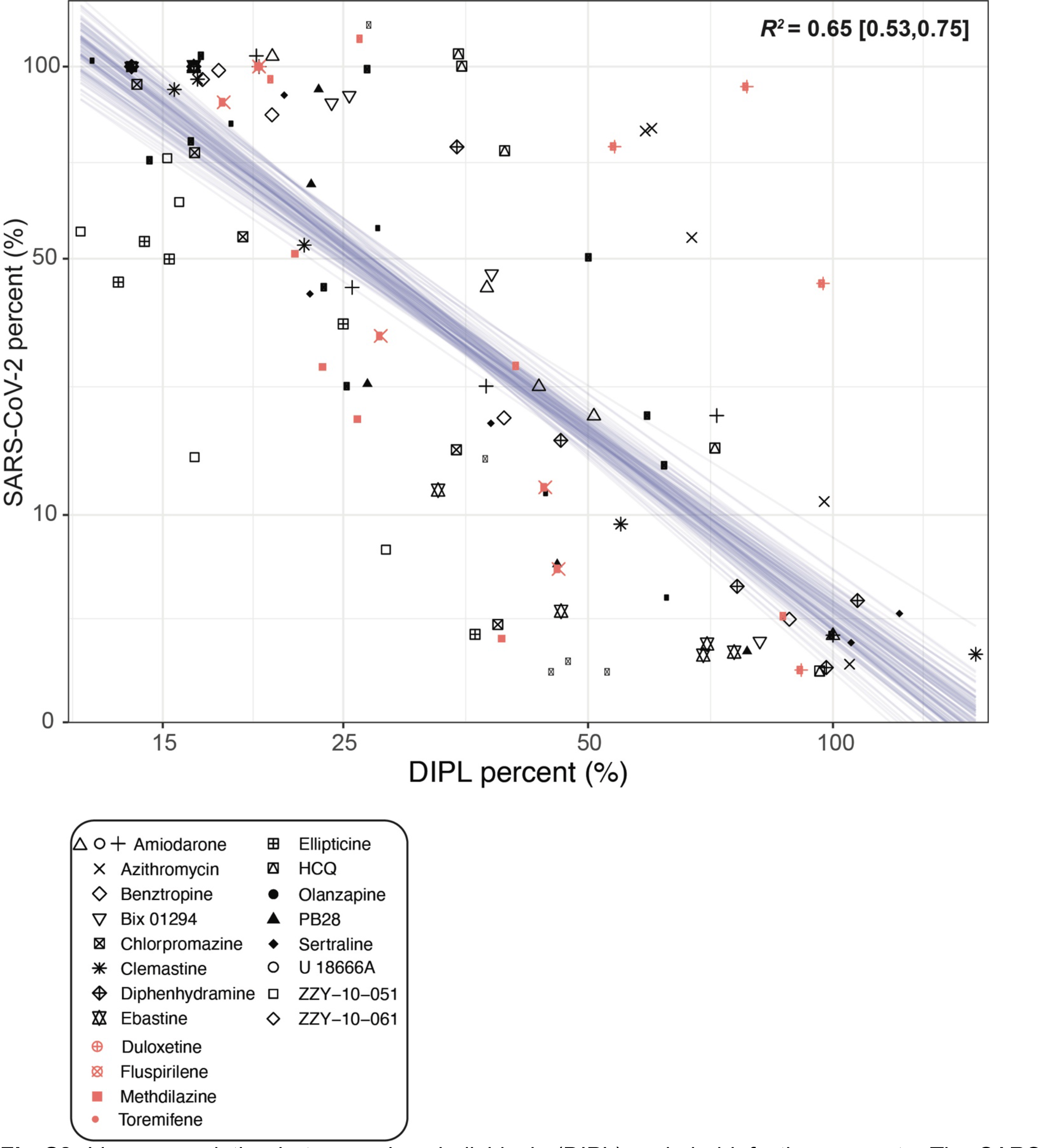
Linear correlation between phospholipidosis (DIPL) and viral infection amounts. The SARS-CoV-2 viral loads and induced phospholipidosis magnitude for each compound and dose are plotted as sqrt(infectivity_mean) ∼ 10*inv_logit(hill*4/10*(log(DIPL_mean)-logIC50). Forty draws from the fit model are shown as blue lines. Salmon points overlaid with the model represent predicted phospholipidosis inducers from the literature (Fig. 5).

**Fig S9.**
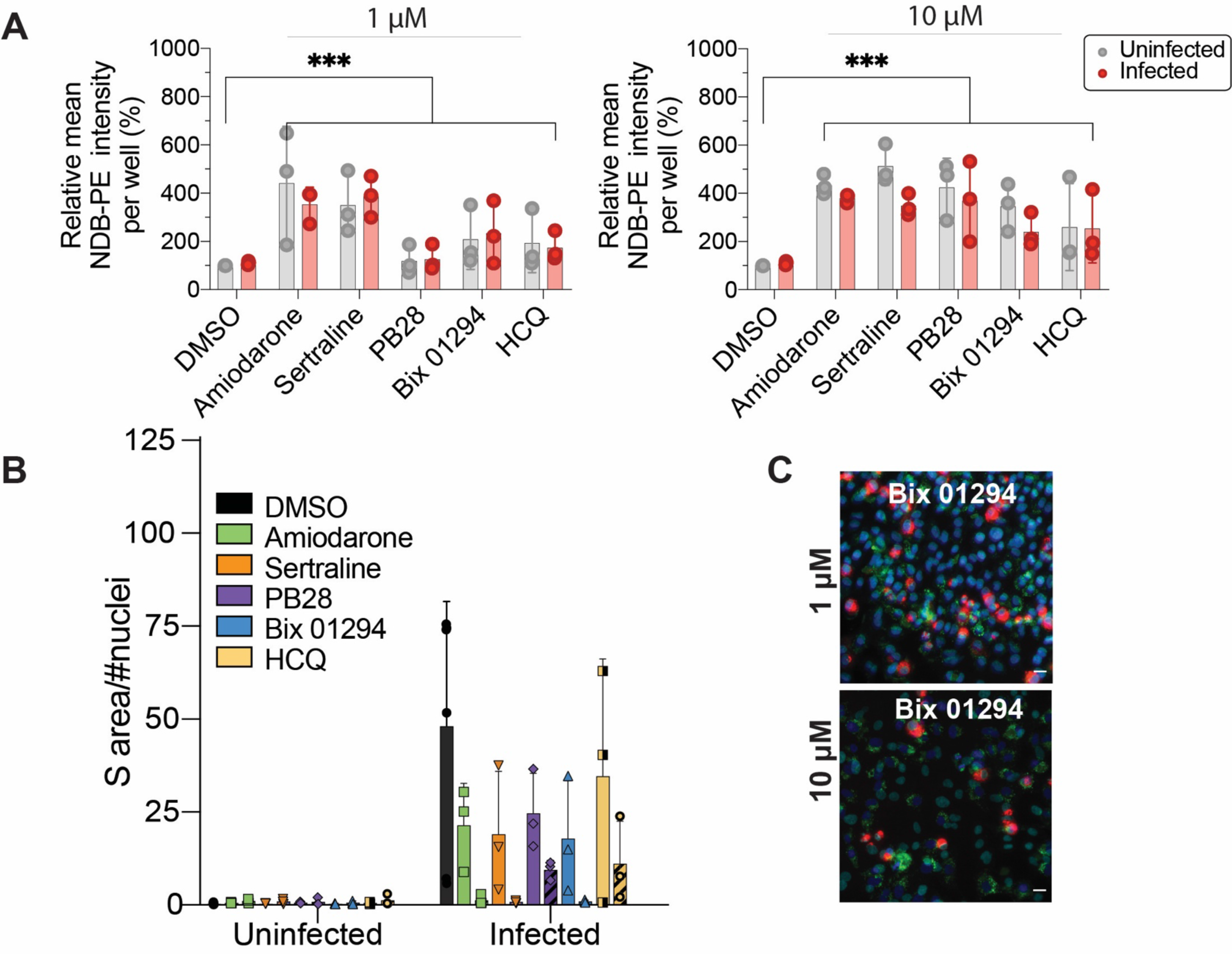
Quantification of phospholipidosis and spike protein in the same cells. **A.** Relative mean ± SD NBD-PE intensity per well percent for 5 molecules and DMSO (1 and 10 µM) in uninfected and SARS-CoV-2 infected A549-ACE2 cells. Data shown from four technical replicates and three biological replicates. Two-way ANOVA main effect of drug treatment, ***P < 0.001. **B.** Spike protein quantification in the same experiment as **A.** for both uninfected and SARS-CoV-2 infected cells, and 1 (solid color bars) and 10 µM (hatched bars) drug treatments. Data represent mean ± SD from four technical and three biological replicates each. Spike protein was quantified as S area / # nuclei per well. **C.** Example images from the costaining experiment measuring phospholipidosis and SARS-CoV-2 Spike protein in infected and uninfected A549-ACE2 cells. Bix 01294 (1 and 10 µM) is shown. Blue = Hoechst nuclei staining; Red = Spike protein staining; Green = NBD-PE phospholipid staining; Yellow = coexpression of spike protein and NBD-PE. Scale bar = 20 µm.

**Fig. S10.**
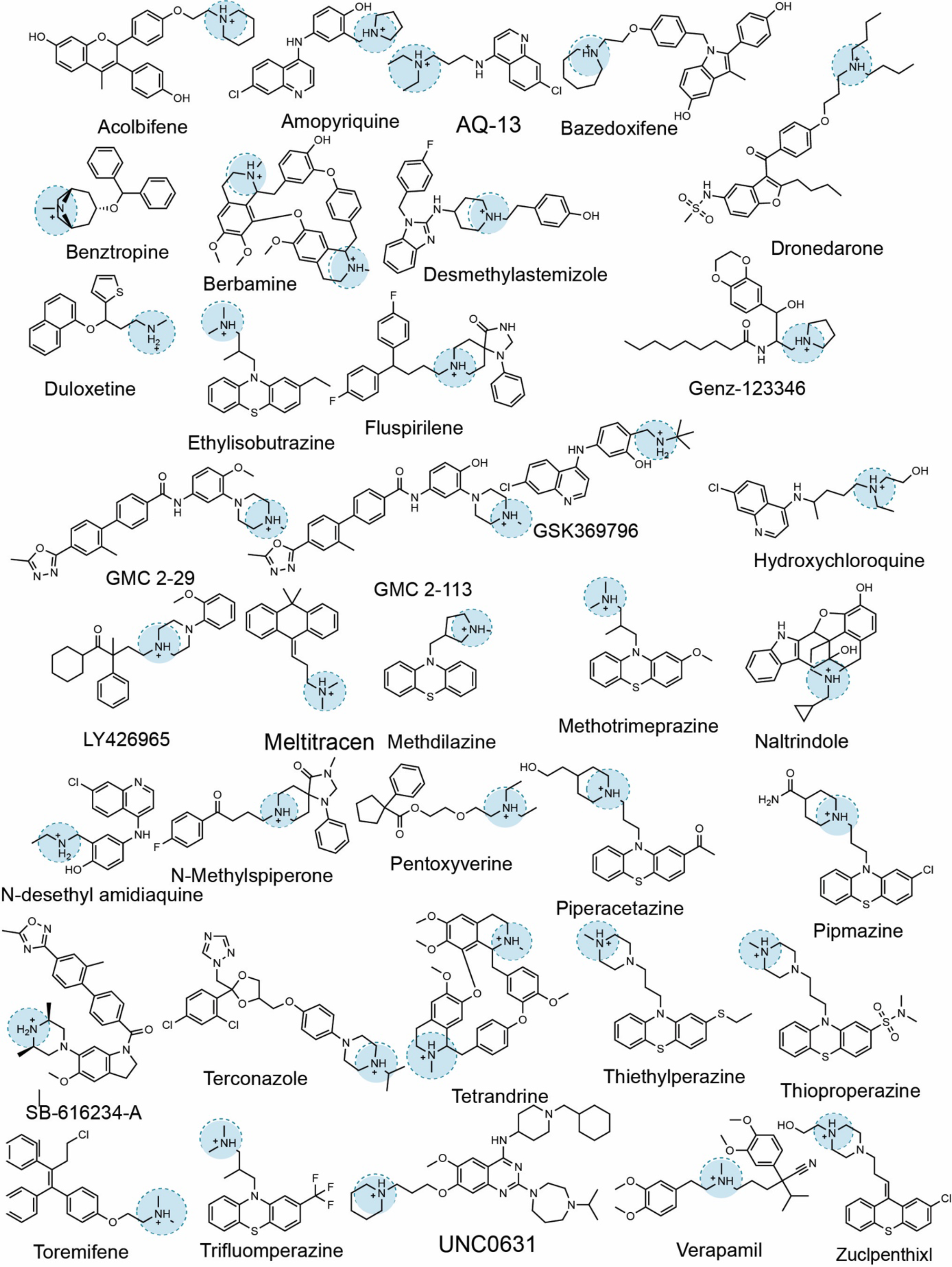
Example cationic amphiphilic drugs identified from SARS-CoV-2 drug repurposing literature predicted to induce phospholipidosis. These molecules have Tanimoto coefficients to known phospholipidosis inducers less than 0.4, and are therefore predicted to induce phospholipidosis.

**Table S3.**
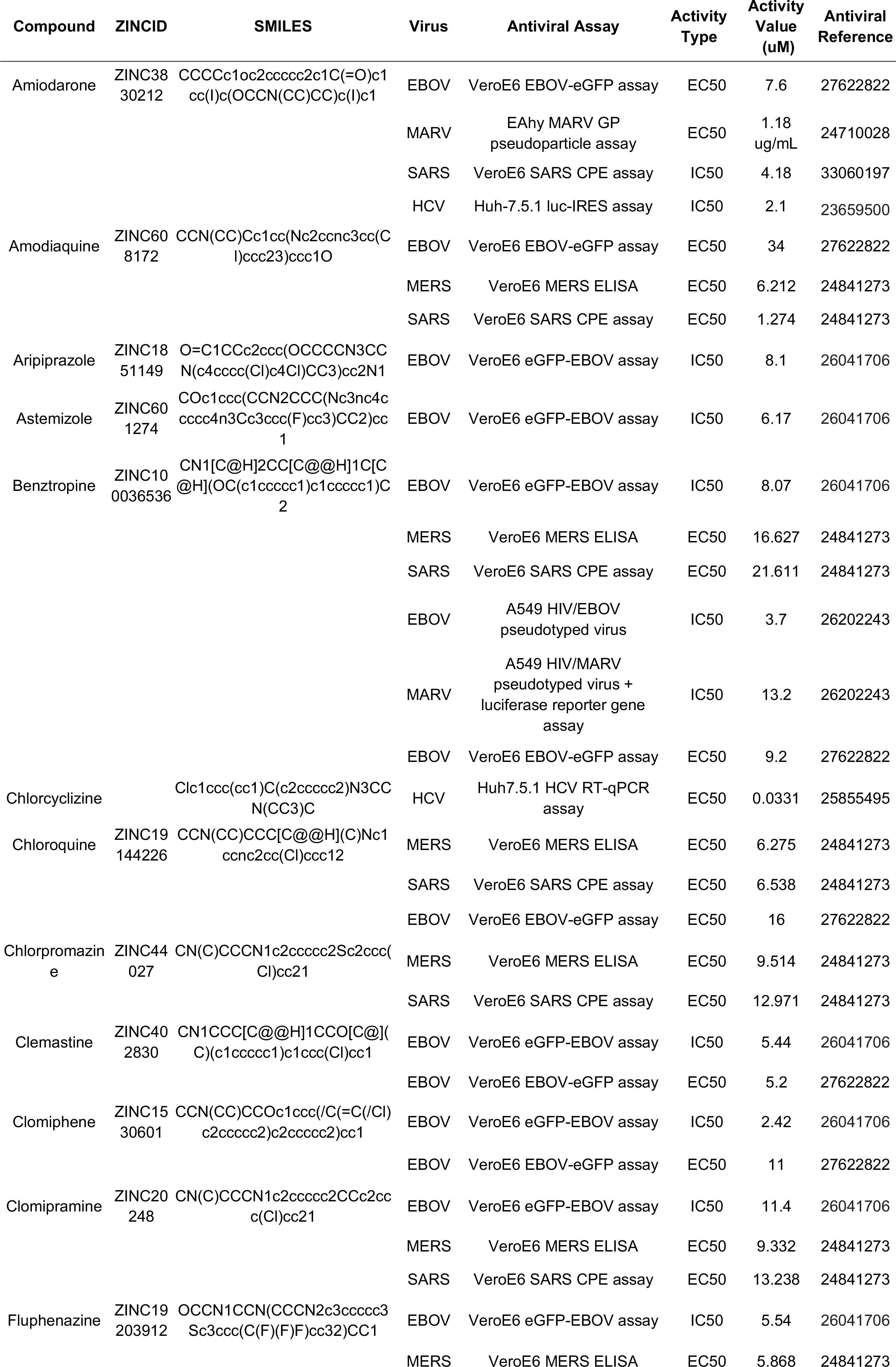
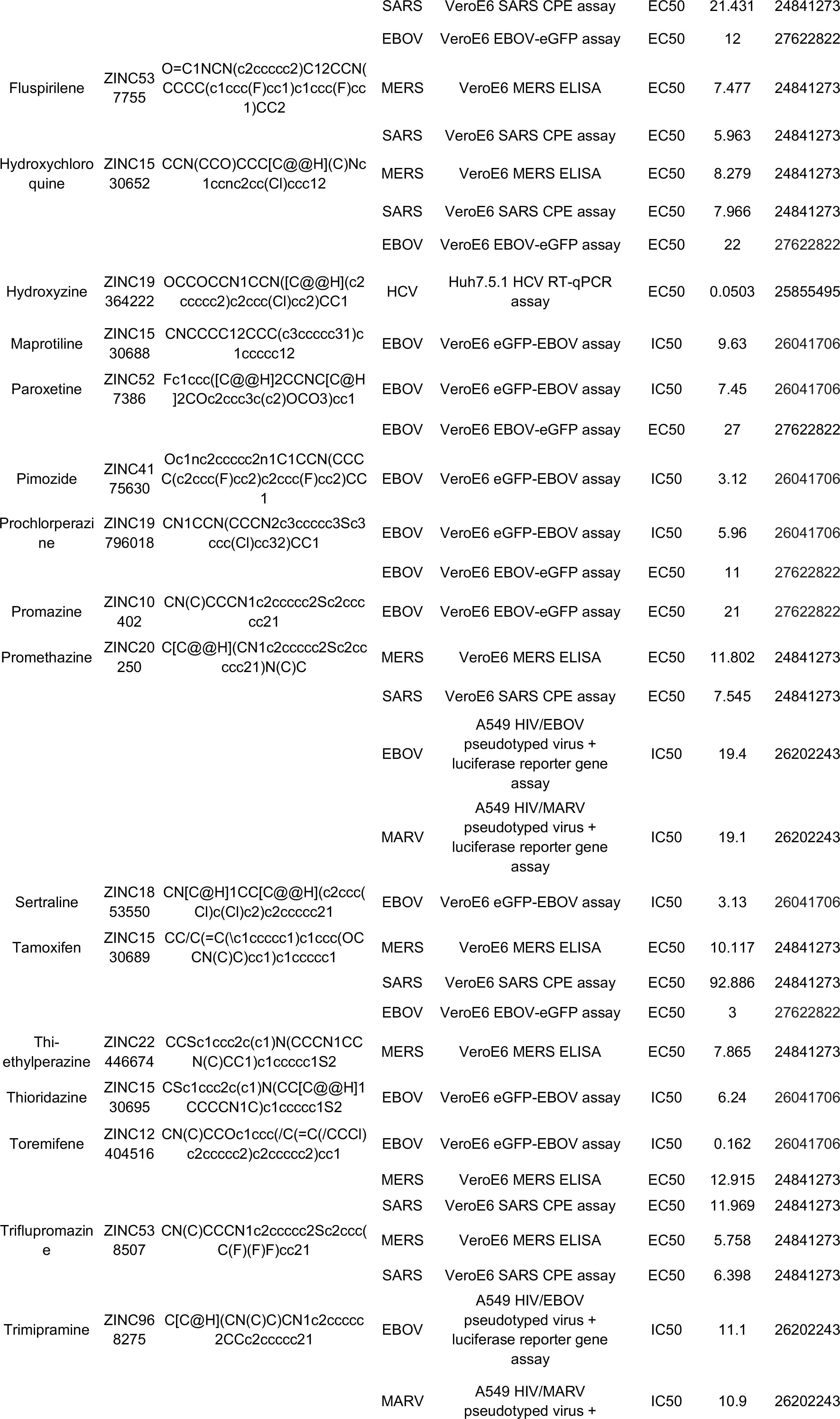
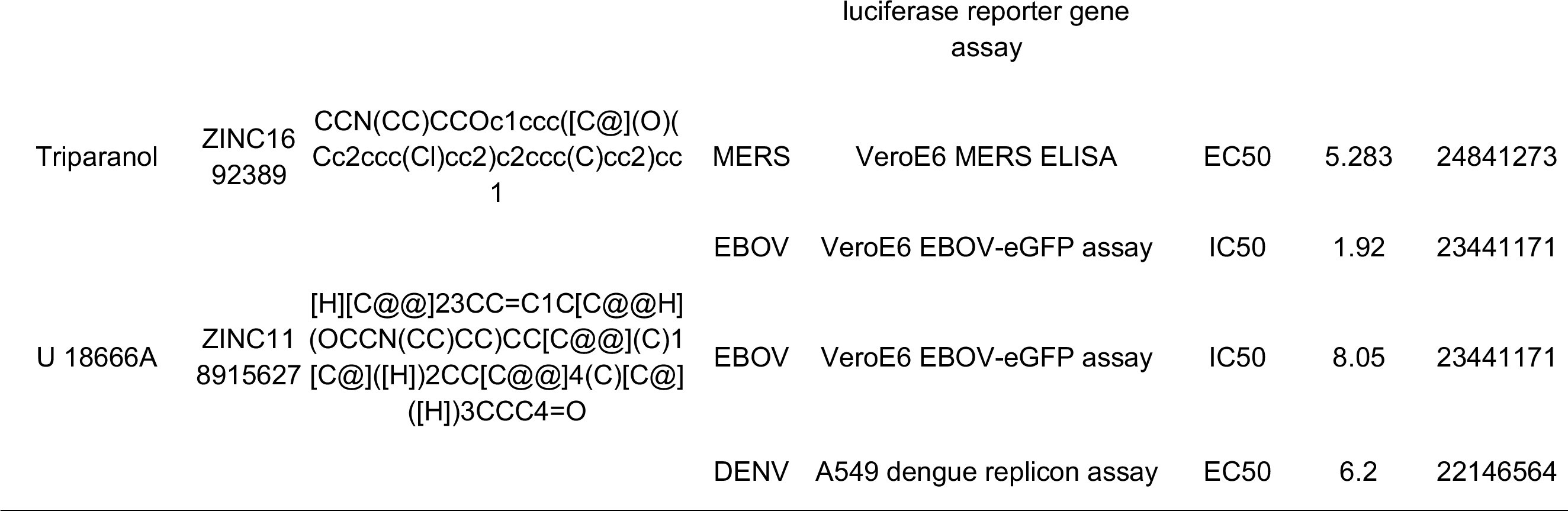
Cationic amphiphilic drugs found active against other viruses in the literature.

**Table S4.** Measured pharmacokinetic parameters for Amiodarone.

| Amiodarone |  | Dose<br>Route | Dose<br>(mg/kg) | T<sub>max</sub><br>(min) | C<sub>max</sub>,<br>ng/mL<br>(g) | AUC<sub>0→tlast</sub><br>ng*min/mL<br>(g) | AUC<sub>0→∞</sub><br>ng*min/<br>mL (g) | T<sub>1/2</sub><br>(min) | K<sub>el</sub><br>(min<sup>-1</sup>) | C<sub>max</sub><br>(nM) |
| --- | --- | --- | --- | --- | --- | --- | --- | --- | --- | --- |
| MWT (Da)<br>681.8 | Plasma | i.p.<sup>a</sup> | 0.3 | 30 | 37.4 | 18800 | 34200 | 1270 | 0.000548 | 55 |
|  |  | i.p. | 1 | 30 | 73.5 | 42000 | 52500 | 583 | 0.00119 | 108 |
|  |  | i.p. | 3 | 30 | 178 | 96700 | 144000 | 769 | 0.000901 | 261 |
|  |  | i.p. | 10 | 15 | 519 | 205000 | 278000 | 585 | 0.0118 | 761 |
|  |  | i.p. | 30 | 60 | 1070 | 476000 | 577000 | 505 | 0.00137 | 1569 |
|  | Lung | i.p. | 0.3 | 360 | 120 | 121000 | 162000 | 698 | 0.000993 | 176 |
|  |  | i.p. | 1 | 120 | 328 | 310000 | 375000 | 599 | 0.00116 | 481 |
|  |  | i.p. | 3 | 120 | 1240 | 1110000 | 1230000 | 501 | 0.00138 | 1819 |
|  |  | i.p. | 10 | 120 | 8020 | 3650000 | 4870000 | 645 | 0.00107 | 11763 |
|  |  | i.p. | 30 | 120 | 13900 | 8930000 | 17900000 | 1250 | 0.000554 | 20387 |

<sup>a</sup>i.p. = intraperitoneally

**Table S5.**
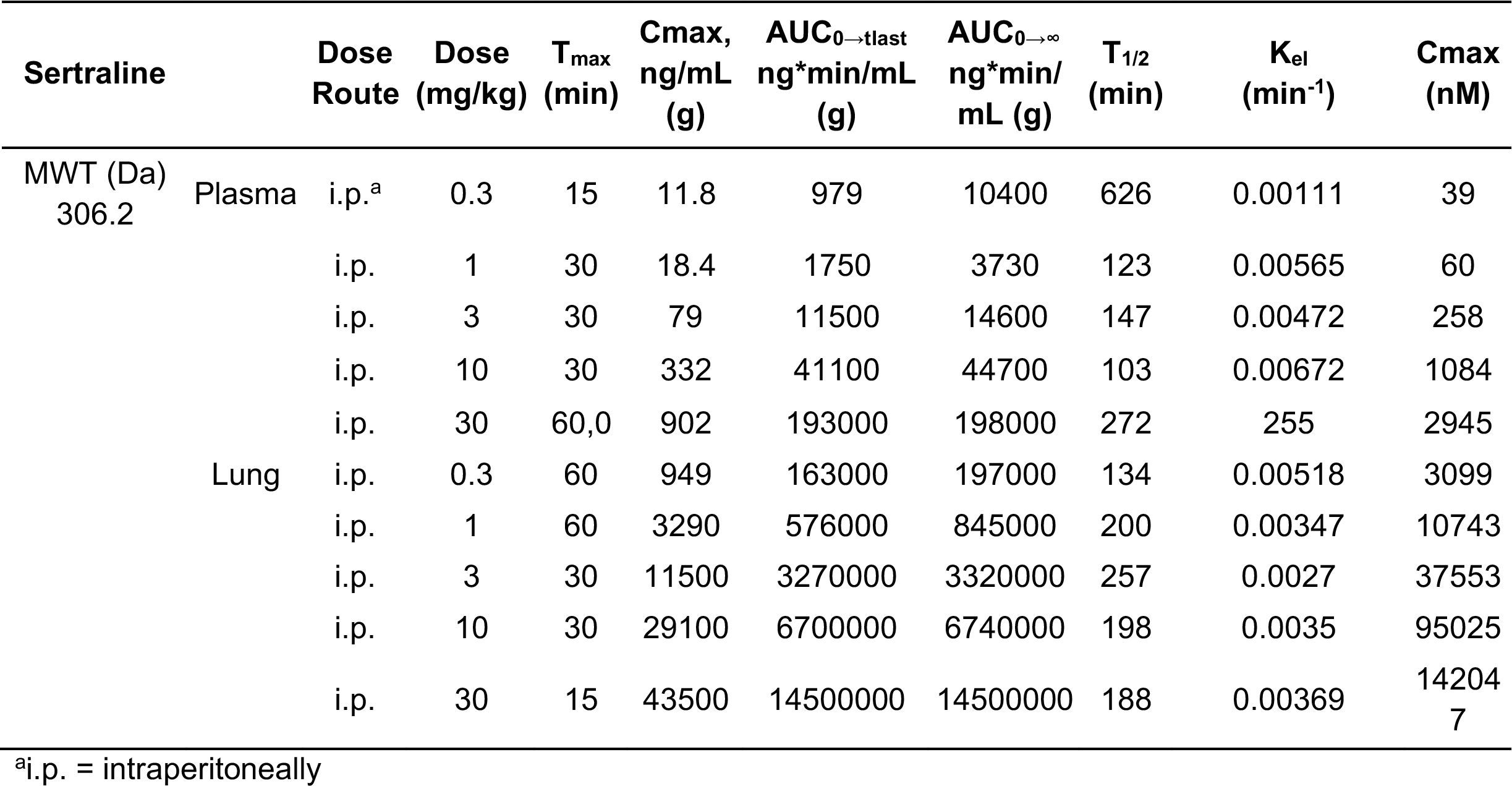
Measured pharmacokinetic parameters for Sertraline.

**Table S6.**
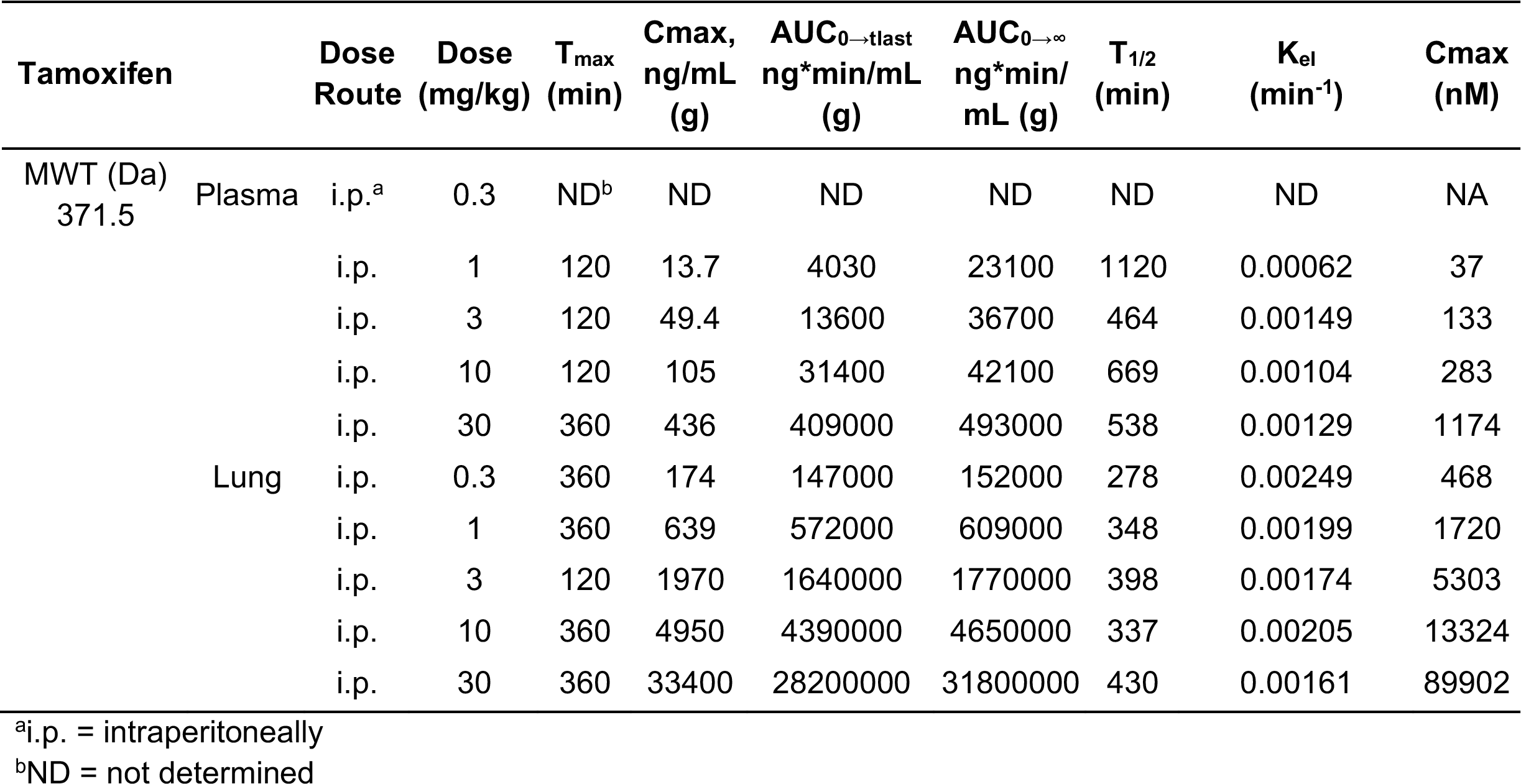
Measured pharmacokinetic parameters for Tamoxifen.

**Table S7.**
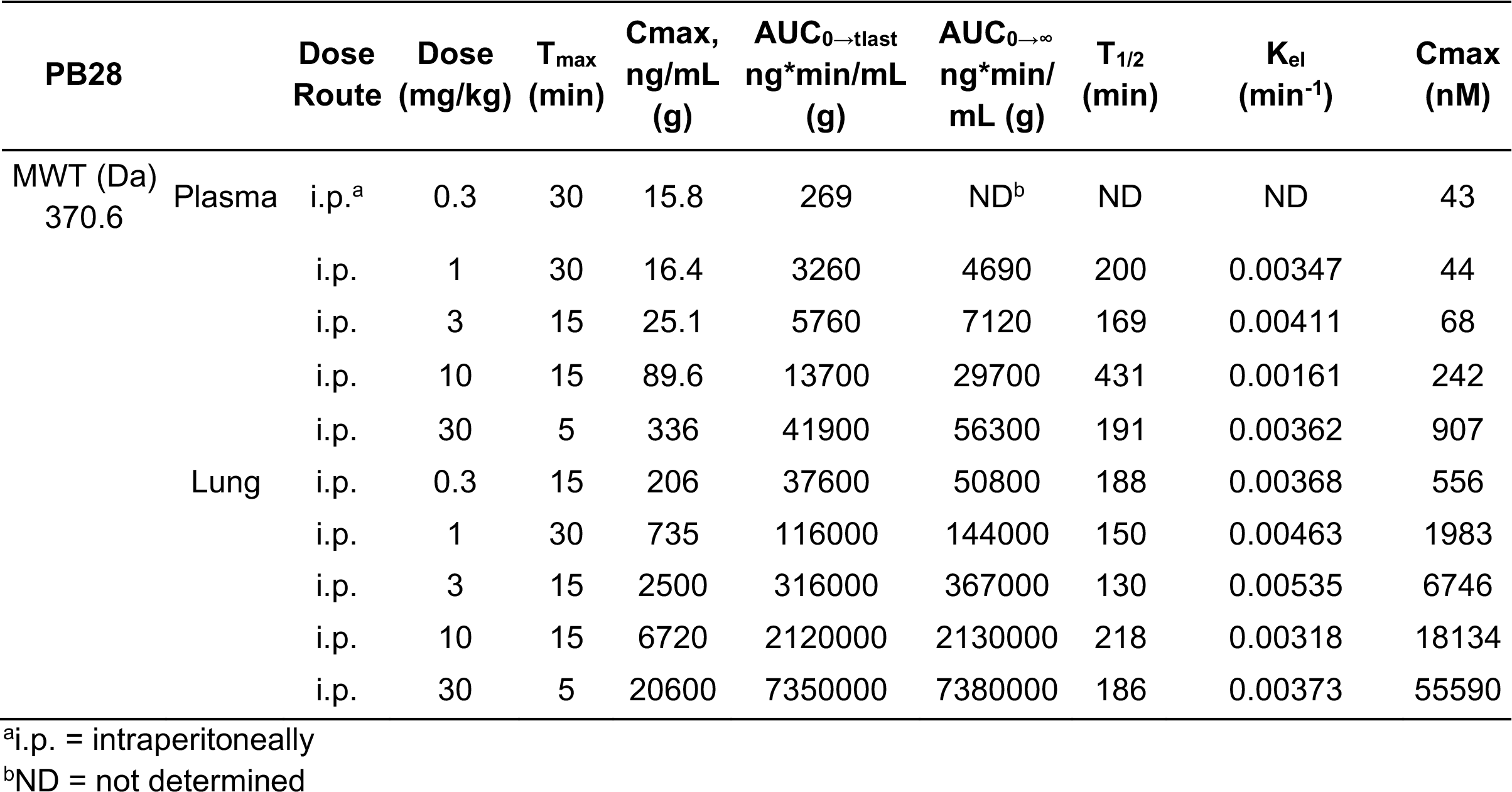
Measured pharmacokinetic parameters for PB28.

**Table S8.**
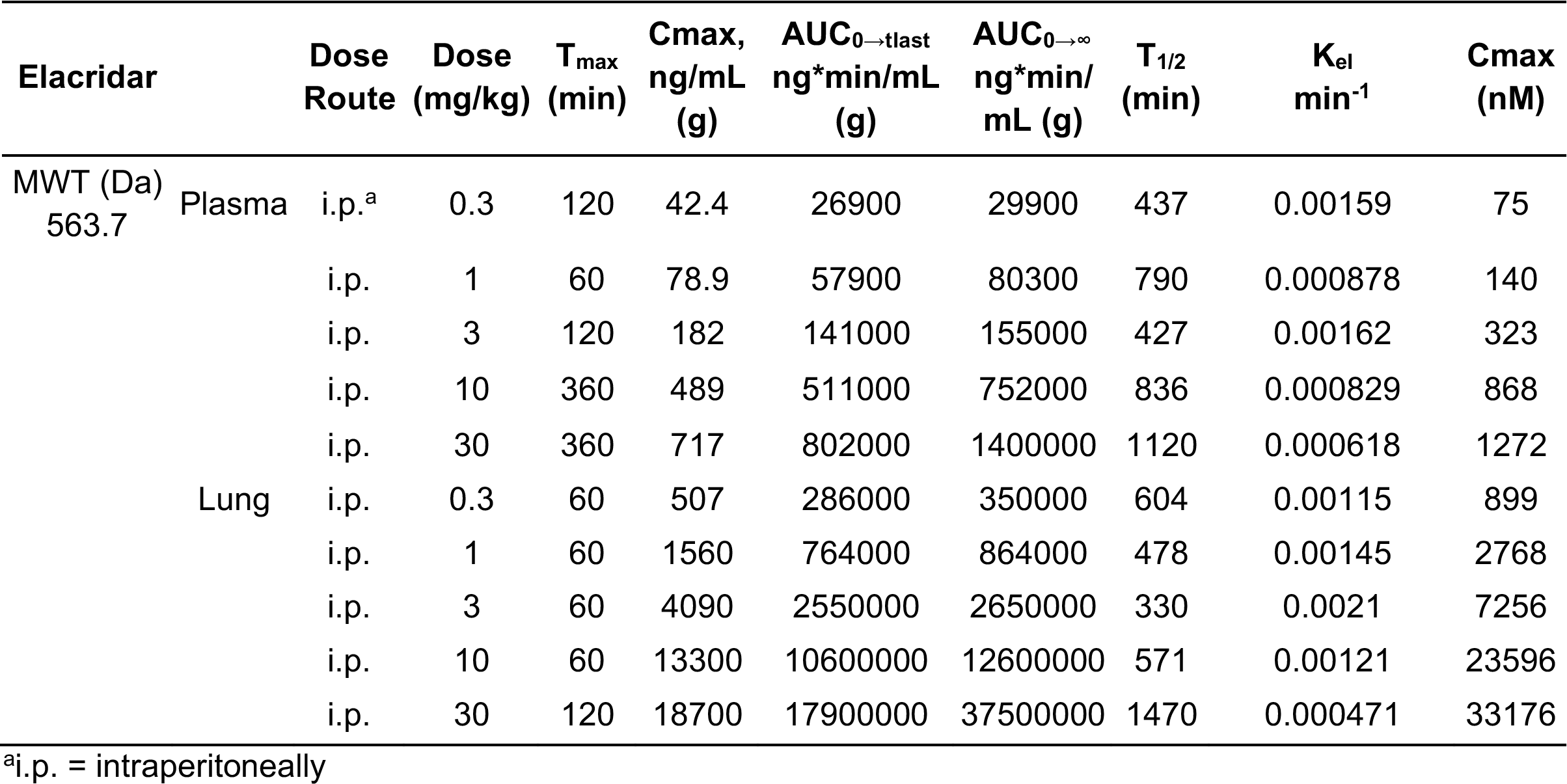
Measured pharmacokinetic parameters for Elacridar.

**Table S9.**
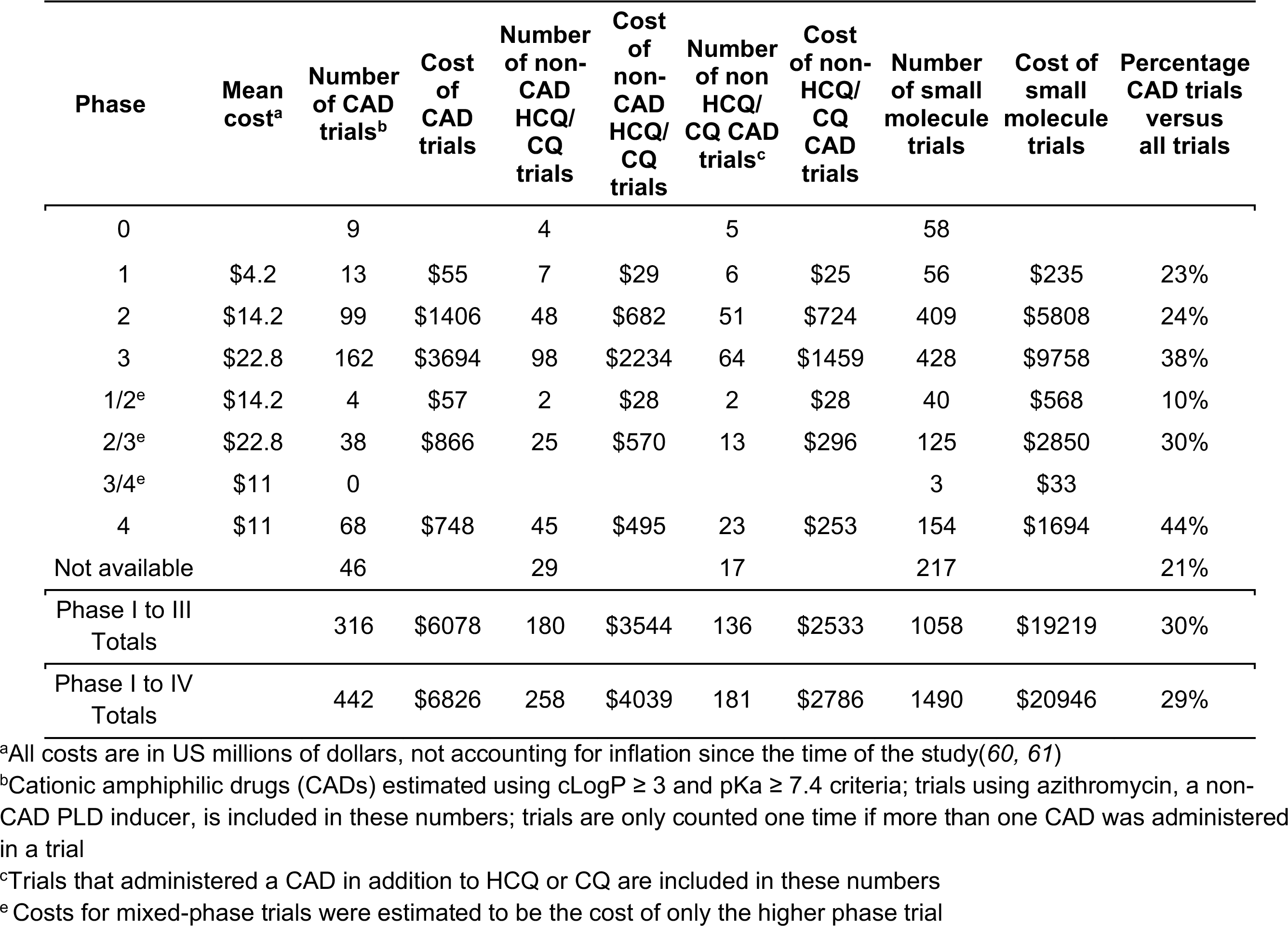
Estimates of expenditures of COVID-19 cationic amphiphilic drug clinical trials

